# Large-scale Signal Propagation Modes in the Human Brain

**DOI:** 10.1101/2024.11.22.624801

**Authors:** Youngjo Song, Pyeong Soo Kim, Benjamin A. Philip, Taewon Kim

## Abstract

The brain’s large-scale temporal dynamics play a crucial role in understanding its operations, but developing a cohesive framework to integrate the potentially extensive array of spatiotemporal patterns remains elusive. Our work addresses this gap by identifying multiple large-scale signal propagation modes in resting-state fMRI time series under a unified methodological framework. We found five distinct modes that effectively predict future blood-oxygen-level-dependent (BOLD) signal dynamics, each reconciling transitions between well-known large-scale brain networks into coherent spatiotemporal units. By utilizing these coherent units, our approach circumvents the need to explore combinatorial explosion of transitions between potential states, enabling parsimonious modeling and effective prediction of whole-brain temporal evolution. Each mode captures specific operational dimensions of neural resource allocation, ensuring their interpretability. Importantly, we showed that complex spatiotemporal features emerge from the superposition of these few propagation modes, unifying a broad spectrum of well-known brain dynamics phenomena. Our results lay the groundwork for a unified framework to understand large-scale spatiotemporal brain organization. Moreover, individual differences in mode expression profiles correlate with general cognitive abilities, exhibit heritability, and demonstrate cross-task stability, underscoring their functional significance. This could lead to efficient methods for characterizing functional fingerprints and advancing diagnostic approaches for neurological disorders.

## Article

The large-scale neural activity can be summarized to a finite set of recurring spatial patterns of activity or connectivity [e.g., 1, 2, 3], commonly referred to as “brain states.” These brain states have been successfully employed as fundamental units for bridging functional organization with cognitive processes, integrating observations across species and scales [4]. However, the brain is known to exhibit fluid reconfiguration [5–7], frequently unfolding through structured temporal sequences of spatial patterns aligned with distinct functional or cognitive processes [8–10]. Such dynamics raise questions about the current practice of relying on static patterns as the primary unit for understanding brain function.

A prototypical illustration is bidirectional signal propagation along the cortical hierarchy, documented across species [11–13]. This propagation demonstrates ordered temporal progression along the axis spanning unimodal regions (e.g., primary sensory/motor areas) and transmodal regions (e.g., the default mode network, DMN). While this axis (i.e., cortical hierarchy) is recognized by static contrasts (e.g., connectivity gradients, cytoarchitectonic differences [14–16]) and aligns with previously recognized spatial patterns (i.e., unimodal vs. DMN [1, 17]) at its extremes, its functional dimension is more explicitly reflected in this temporal progression: bottom-up signals originating in unimodal regions integrate sensory inputs, while top-down signals from transmodal regions modulate predictions and attentional processes [18–22].

A related example is the dynamic reallocation of resources during tasks, where activity shift from the task-negative (TN) network (i.e., DMN) to the task-positive (TP) network (e.g., central executive network, CEN) [23–28]. Although TN and TP networks are robustly anti-correlated [29] and have been traditionally regarded as opposing systems (i.e., internally oriented vs. externally engaged cognition), emerging evidence suggests a more integrated relationship. First, a single spatial pattern typically lasts a few seconds—too brief to solely account for such sustained a cognitive state [2, 10, 30]. Second, the quasi-periodic oscillations between TN and TP reliably occurs during both rest and task conditions [31, 32], demonstrating that these states are not strictly tied to specific contexts (e.g., rest vs. task). Instead, they reflect a cooperative rhythm underlying cognition [33, 34]. These insights suggest a new perspective on the classical “triple network” hypothesis [35], which proposes that the salience network (SN) orchestrates TN–TP transitions [36–38] by detecting behaviorally relevant stimuli [39]. In light of these dynamic interplays, it now appears more accurate to characterize the SN’s role as modulating ongoing temporal patterns [e.g., 31, 32] rather than simply gating static network configurations.

Overall, these insights underscore the necessity of conceptualizing brain function in terms of evolving spatiotemporal dynamics [for review, see ref. 9]. Indeed, growing evidence suggests that examining temporally evolving patterns can better reflect cognitive traits and yield more sensitive biomarkers [40–43]. However, integrating the temporal dimension into coherent units poses significant challenges. First, while the brain’s flexible reconfiguration implies that all sequential combinations of brain states (i.e., spatial patterns) warrant investigation [9], the resulting combinatorial explosion threatens computational feasibility. Second, methodological disparities in identifying spatiotemporal patterns—such as divergent assumptions about *states* (e.g., based on activation vs. connectivity) and *transitions* (e.g., probabilistic vs. deterministic frameworks) [e.g., 2, 10, 11, 12, 13, 30–32, 37]—hinder the integration. Thus, incorporating temporal dimension faces a critical paradox: developing frameworks capable of capturing the richness of sequential brain dynamics while remaining robust to method-dependent and context-dependent variations.

In this context, dynamic mode decomposition (DMD) emerges as a powerful, equation-free, data-driven solution that parsimoniously captures the spatially distributed temporal dynamics in high-dimensional systems [44]. DMD operates by solving eigenvector problems based on a linear approximation of the causal dynamics that govern signal flow within the system (***x***(*t*) = *A****x***(*t* − 1)). This approach decomposes multi-dimensional time-series data into a set of spatiotemporal modes that encapsulate the system’s dominant oscillatory and transitional behaviors, referred to as dynamic modes (DM). (Hereafter, *DM* refers exclusively to dynamic mode, while *DMN* denotes the default mode network.) Unlike traditional methods such as PCA or ICA (which isolate static spatial or temporal covariation) and approaches like co-activation pattern analysis or hidden Markov model (which detect recurring spatial states), DMD isolates *temporally evolving patterns* that explicitly model phase-locked oscillatory dynamics (which can capture signal propagation). Therefore, this approach unifies spatial patterns, temporal evolution, and causal interactions into a single interpretable spatiotemporal unit (i.e., DM). Furthermore, DMD quantifies mode-specific properties (or metrics) such as amplitudes (i.e., engagement level), decay/growth rates (i.e., persistence rate), and oscillation/propagation speed (i.e., progression rate) (hereafter, collectively termed DM metrics), offering a principled way to balance cross-subject commonalities and individual variability in spatiotemporal structure. Beyond descriptive analysis, these mode properties not only allow for system stability assessments and future-state predictions [45–48], but also enable to reconstruct rich sequential dynamics by superposition of a limited set of modes.

In this study, we aimed to address the aforementioned paradox of modeling the brain’s temporal dynamics by systematically employing DMD on whole-brain fMRI time-series data. This framework enabled the identification of five distinct modes of signal propagation within the brain, which outperform existing machine learning benchmarks in predicting future BOLD signals. Superposition of these modes accounts for canonical dynamic features observed in resting-state fMRI. These features encompass the topography of functional connectivity (FC) and its temporal fluctuations, known as time-varying FC (TVFC); FC-based functional gradients [14, 49]; wave propagation along the cortical hierarchy [11–13]; global signals [50]; time-lag structures [51]; quasi-periodic patterns (QPP) [31, 32]; the anticorrelation between TN and TP networks [29]; and hemispheric lateralization observed in resting-state BOLD dynamics [52]. By integrating these dynamics into a single parsimonious model, we demonstrate that seemingly disparate dynamics features emerge as natural consequences of coherent, system-wide causal interactions. The identified DMs correspond to distinct aspects of neural resource allocation, integrating various interactions between major network states (e.g., unimodal networks, DMN, CEN, and SN) into specific functional dimensions.

## Results

To model the temporal dynamics of the global BOLD signal, we applied DMD to resting-state fMRI scans from the Human Connectome Project (HCP) Young Adult S1200 dataset. We utilized the Glasser parcellation [53] to divide the cortical regions into 360 parcels and the Cole-Anticevic Brain-wide network partition [54] to divide the subcortical and cerebellar regions into 356 parcels, resulting in a total of 716 regions of interest (ROIs) (see Methods). Subsequently, we utilized a forward-backward DMD algorithm [55] to compute the DMs of the group-level causal connectivity matrix (i.e., matrix A), which best approximates the evolution of the BOLD signal between adjacent time points across all subjects (see Methods section “Group-level DMD” for details). To ensure physiological relevance, we retained only DMs with oscillation frequencies within the range of 0.01 Hz to 0.1 Hz, consistent with the filter frequency used in fMRI preprocessing.

To determine the optimal number of DMs for accurately describing brain-wide BOLD signal dynamics, we utilized a 5-fold cross-validation approach, evaluating predictive performance for future BOLD signals across varying numbers of DMs. In each fold, the training set was used to estimate the DMs (using the group-level DMD). Each resting-state fMRI run in the test set was further divided into test-fitting and test-validation data. The test-fitting data were used to estimate the DM metrics—parameters that determine how signals sustain their influence and how quickly they propagate within each DM—enabling prediction of the brain-wide signal at subsequent time points. We estimated these DM metrics by projecting the brain-wide BOLD signal onto the DMs and then optimizing to minimize the prediction error 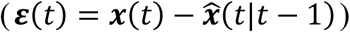,where ***x***(*t*) is the BOLD signal at time *t*. 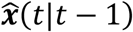 is the predicted BOLD signal at time t based on prior information at time *t* − 1, which was obtained by superposition of the temporal evolution of each DM (for details see subsequent results and Methods section “Subject-level estimation of DM metrics”). For model comparison, we utilized the cross-validated *R*^2^ measure, which quantifies the proportion of variance in the signal that the model can explain based on the prediction error within the test-validation segment (see Methods section “Optimizing the number of DMs”). The methodology for DMD and DM metric estimation is illustrated in Figure 1a.

**Figure 1.**
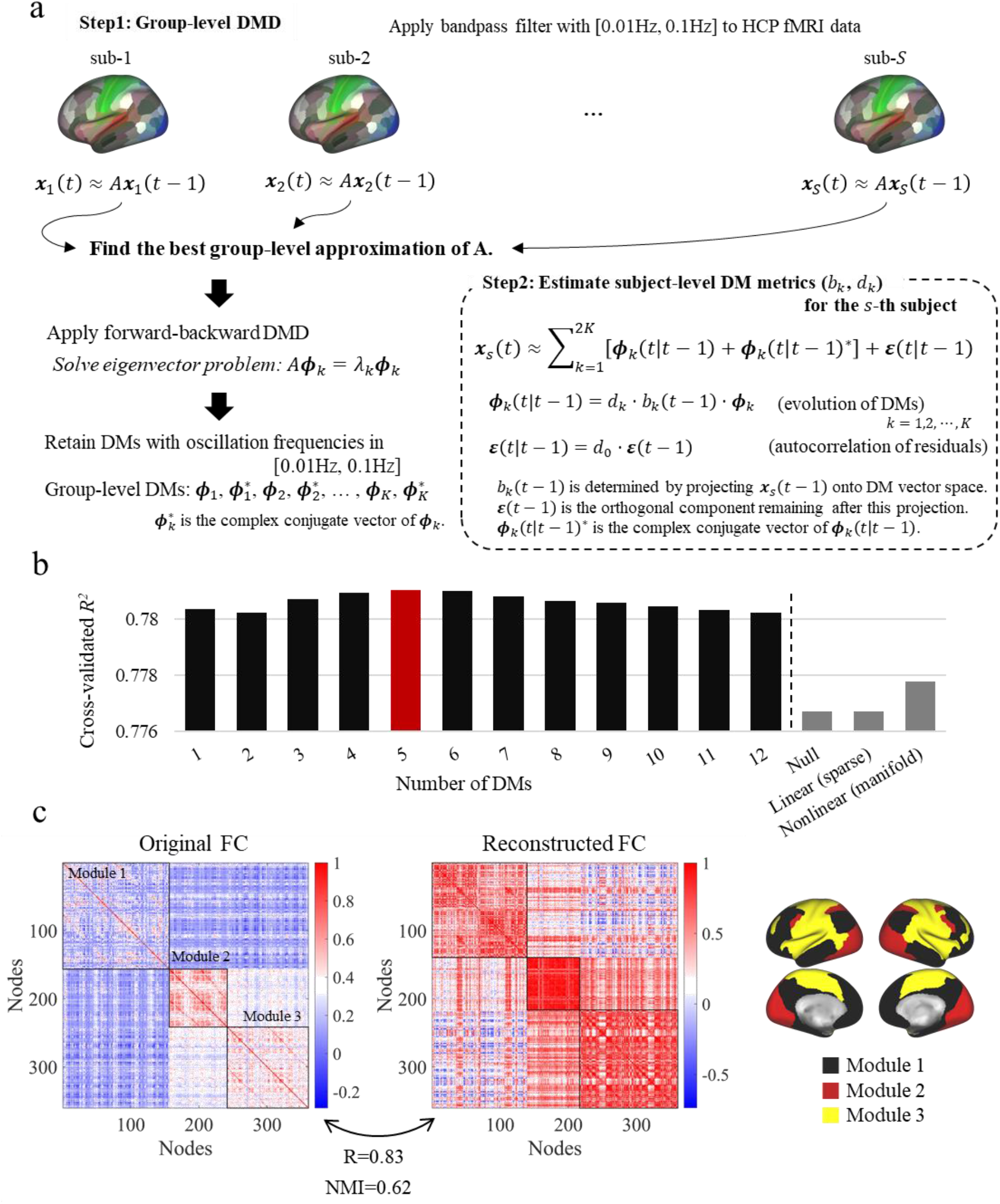
**(a)** Schematic of group-level Dynamic Mode Decomposition (DMD) applied to resting-state fMRI data and the estimation of subject-specific DM metrics with *K* DMs: mode-engagement level (|*b*_*k*_|), persistence rate (|*d*_*k*_|),), and progression rate (tan^−1^(Re(*d*_*k*_) / Im(*d*_*k*_))). **(b)** Prediction performance, indicated by cross-validated *R*^2^, using different numbers of DMs, as well as using alternative models (i.e., LASSO and manifold-based model) for comparison. DMs were incrementally included based on the magnitude of their group-level evolution coefficients (*λ*_*k*_; see Methods), reflecting their contributions to the overall dynamics. Models incorporating DMs consistently outperformed the best previously evaluated models in predicting future BOLD signals [56]. The highest prediction accuracy was achieved using five DMs. **(c)** FC network reconstructed by the five DMs. **Left panel:** FC matrix of cortical BOLD time courses (i.e., original FC matrix). **Middle panel:** Correlation matrix reconstructed from the five DMs. **Right panel:** Module assignments for each ROIs using the cortical FC matrix. Both the original and reconstructed correlation matrices are ordered and outlined according to the modular structure identified by Newman’s spectral community detection algorithm, respectively. The community analysis revealed three modules corresponding to the DMN, visual, and SM networks. Although the reconstructed FCs are predominantly positive, the correlation patterns between the original and reconstructed matrices are highly similar (R = 0.83). Additionally, the modular structures of the original and reconstructed matrices show strong similarity (NMI = 0.62), with 87% of ROIs correctly matched.

Remarkably, the results of DMD (i.e., DMs) remained highly consistent across all five cross-validation folds (Supplementary Result 1), with each fold identifying twelve DMs, demonstrating the robustness of our DMD approach. Cross-validation results indicated that five DMs were optimal to predict the temporal evolution of BOLD signal (Figure 1b). Notably, even when using a single DM, we achieved a lower prediction error (i.e., higher cross-validated R^2^) compared to the linear and non-linear models, which had previously demonstrated the best performance in previous evaluations [56].

The reconstructed cortical FC matrix using these five DMs showed a strong correlation with the original group-FC matrix (R = 0.83; Figure 1c). Network community analysis revealed a strong correspondence between the community structures of the original and reconstructed FC networks, with a normalized mutual information (NMI) value of 0.62—indicating that 87.5% of nodes are matched, thus reflecting high similarity in community organization. Furthermore, the TVFC, summarized by standard deviation (std), also showed a strong correlation between the original and reconstructed TVFC matrices (R = 0.75; Supplementary Figure 3a), suggesting that temporal fluctuations of FC are captured by the temporal evolution of the five DMs. These findings collectively suggest that the five DMs parsimoniously encapsulate the essential dynamics and organizational features of brain-wide BOLD signal activity, effectively reconstructing both the static and dynamic aspects of FC. Importantly, these five DMs correspond to temporal transitions between well-recognized brain-wide networks, which will be elaborated in subsequent sections.

### DM1: Principal DM accounts for signal propagation along the cortical hierarchy

The temporal dynamics of a linear system are predominantly governed by the DM with the largest eigenvalue [57]. In our analysis of resting-state fMRI data, the DM with the highest eigenvalue—referred to as the “principal DM”—demonstrated signal propagation between unimodal regions (i.e., primary visual, secondary visual, primary SM, and primary auditory areas) and the transmodal areas (largely overlapping with the DMN) (Figure 2a). Consistent with its largest eigenvalue, this principal DM emerged as the most actively engaged in the fMRI scan (see the left panel of Supplementary Figure 10a; later sections detail the methods for quantifying engagement level of DM). Importantly, in our cross-validation analyses aimed at predicting future BOLD signals (Figure 1b), the utilization of solely the principal DM resulted in superior predictive performance when compared to both linear and non-linear models. Moreover, this principal DM accounted for the majority of variance explained across all predictions, irrespective of the number of DMs used.

**Figure 2.**
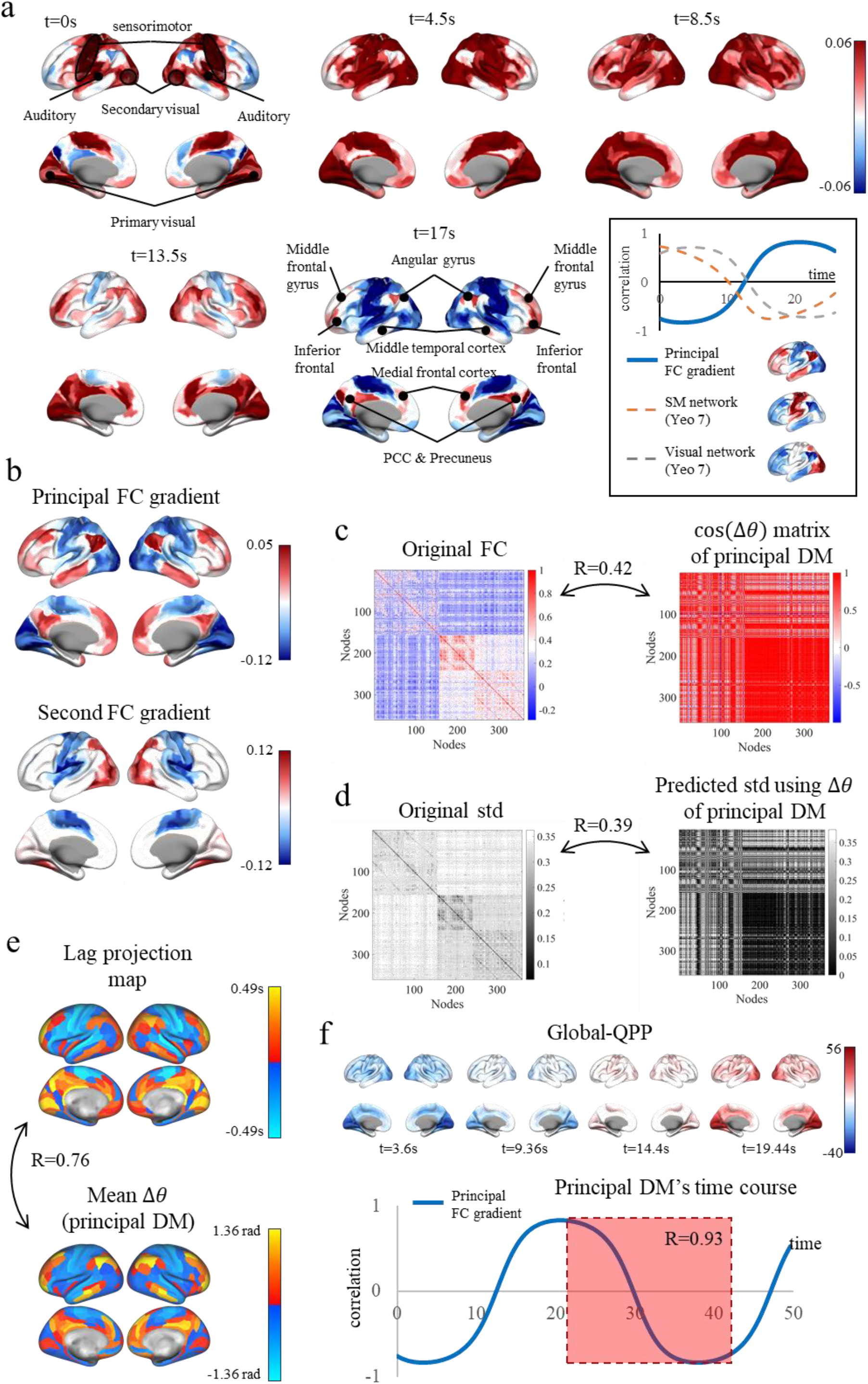
**(a)** Reconstructed timepoints of the principal DM. In this illustration, unimodal regions—including the primary visual, secondary visual, auditory, and SM regions—are positively activated at *t* = 0. As time progresses, the signal gradually propagates to transmodal cortical regions that largely overlap with the DMN. This propagation is evidenced by the correlation of spatial pattern at each time point with the principal FC gradient, SM network, and visual network (the latter two networks were obtained from Yeo’s 7-network parcellation; see Methods), as shown in the inset. Near the initial time point (t ≈ 0), the spatial pattern of the principal DM shows a strong correlation with the SM and visual networks (R > 0.7) and a negative correlation with the principal FC gradient. Around t ≈ 17.5, the spatial pattern exhibits a strong correlation with the principal FC gradient (R > 0.7). Note that DMs demonstrate periodic evolution. Consequently, the principal DM also reflects transmodal activation propagating back to unimodal regions, indicating bidirectional signal propagation along the unimodal-transmodal axis. Refer to the lower panel of Figure 2e for reconstructed time points over extended durations, summarized by the correlation with the principal FC gradient. **(b)** Spatial topographies of FC gradients. **Upper panel:** Principal FC gradient (first principal component). **Lower panel:** Second FC gradient (second principal component). **(c) Left panel:** Original FC matrix computed from cortical BOLD time series. **Right panel:** Exact reconstruction of the FC matrix for the principal DM, obtained by calculating the cosine of the phase difference (Δ*θ*) between pairs of ROIs. **(d) Left panel:** Std matrix representing temporal fluctuations of FC in the cortical BOLD time series (i.e., the original std matrix). **Right panel:** Reconstruction of the standard deviation matrix of temporal FC fluctuations using the phase differences between pairs of ROIs in the principal DM. Even using this single DM, the reconstructed FC and std matrices show moderate correlations with the original matrices (R = 0.41 and R = 0.39, respectively). **(e) Upper panel:** Lag projection map derived from cortical BOLD time series, illustrating the average time lag (in seconds) for each cortical ROI. **Lower panel:** Average phase difference map of the principal DM, showing the average phase lag (in radians) for each cortical ROI. The strong correlation between the lag projection and average phase difference maps (R = 0.72) indicates that the time lags in the BOLD signal are predominantly influenced by the principal DM. **(f) Upper panel:** Spatiotemporal pattern observed in the global QPP (i.e., QPP obtained without GSR). The global QPP exhibits a widespread pattern of activity with positive correlations across brain regions. **Lower panel:** Reconstructed time points of the principal DM, summarized by the correlation with the principal FC gradient at each time point. The global QPP showed a strong correlation with interval when the principal DM displays a decrease in correlation with the principal FC gradient (R = 0.93), indicating that the global QPP largely reflects the temporal evolution of the principal DM.

The observed signal propagation between unimodal and transmodal areas is supported by the correlation between the spatial activity patterns of the principal DM at each reconstructed time point and the principal FC gradient (see the upper panel of Figure 2b for topography of the principal FC gradients). This correlation achieved a peak negative correlation of –0.83 during activation of unimodal regions and a peak positive correlation of 0.83 during activation of transmodal regions (see the inset of Figure 2a). The principal FC gradient is well-recognized for its alignment with the cortical hierarchy defined both anatomically and functionally [14]. Thus, these findings suggest that the most dominant brain-wide propagation (i.e., the principal DM) is driven by information exchange along the cortical hierarchy (i.e., unimodal-transmodal axis), aligning with previously observed wave propagation patterns [11–13]. Supplementary Video 1 depicts the signal propagation within the principal DM.

Building on the principal DM’s prominent influence, we hypothesized that the principal DM predominantly governs the key characteristics of resting-state dynamics. Consistent with this hypothesis, the cosine values of phase differences between ROIs—an exact reconstruction of FC between phase-locked ROIs—within the principal DM were moderately correlated with the cortical FC matrix (R = 0.42; Figure 2c). Furthermore, the std of temporal fluctuations in FC, as observed in the fMRI signal, was also moderately correlated with the predicted std calculated using only the phase differences in the principal DM (R = 0.39; Figure 2d). These findings suggest that the global FC distribution and its fluctuations are significantly influenced by signal propagation along the cortical hierarchy.

To further elucidate this relationship, we examined the association between the time-lag structure and the ROI’s phase differences within this DM. Time lag refers to the delay or causal dependence between two brain signals, where one signal is temporally delayed relative to another, thereby representing the overall temporal precedence or lag of each ROI (around ±2 seconds) [51]. Similarly, the phase difference between ROIs quantifies the extent to which one ROI’s activation is temporally delayed relative to another within this DM. We observed a strong correlation between the lag projection—the average time lag between ROIs—calculated with fMRI signal and the average phase difference in the principal DM (R = 0.76; Figure 2e), supporting the major contribution to the brain’s overall dynamics. Additionally, the time-series of the principal DM achieved a peak correlation with QPP [32]—a time-lag phenomenon occurring at longer timescales (~20 seconds)—of R = 0.93 (Figure 2f). This high correlation is noteworthy, given that time-lag structures and QPP can be considered spatiotemporal patterns occurring at different timescales, suggesting that both arise from the brain-wide signal propagation captured by the principal DM. Furthermore, the global signal was predominantly correlated with the global activity of the fMRI signal projected onto the principal DM (mean R = 0.63, the highest among the five DMs; see Supplementary Figure 5), underscoring the principal DM’s dominant contribution to the global signal.

Extending beyond cortical regions, the principal functional gradients of subcortical areas—including the hippocampus, striatum, thalamus, brain stem, and amygdala—and the cerebellum also exhibited strong alignment with the principal DM, each achieving peak correlations exceeding R > 0.7 (see Supplementary Figure 7). These functional gradients capture the dominant variations in functional organization within subcortical and cerebellar FC. Therefore, this observation further highlights the critical role of the principal DM in signal propagation along the dominant functional variations, not only within cortical regions but also robustly across subcortical structures and the cerebellum. Overall, these results establish the principal DM as a key unifying factor in explaining various brain network behaviors.

### DM2&3: Dynamic shift between the three major large-scale networks (DMN, CEN, and SN)

In our analysis, we identified two distinct DMs that were second and third in terms of overall engagement throughout the scanning sessions, each reflecting interactions and transitions among the DMN, CEN, and SN—three major intrinsic large-scale brain networks. These DMs exhibited unique transition pathways contingent upon SN activation (Figure 3). Specifically, one mode involved a transition from the SN to the DMN, accompanied by suppression of the CEN, which we term the “SN-to-DMN” DM (Figure 3a). Conversely, the other mode involved a transition from the SN to the CEN, also accompanied by suppression of the DMN, was referred to as the “SN-to-CEN” DM (Figure 3b). This contrasting dissociation of the DMs highlights the SN’s role as a dynamic switching hub capable of modulating the activity of the DMN and CEN by either facilitating or suppressing them. These findings align with recent perspectives that position the SN as a causal signaling hub responsible for network switching between the DMN and CEN [35, 58, 59].

**Figure 3.**
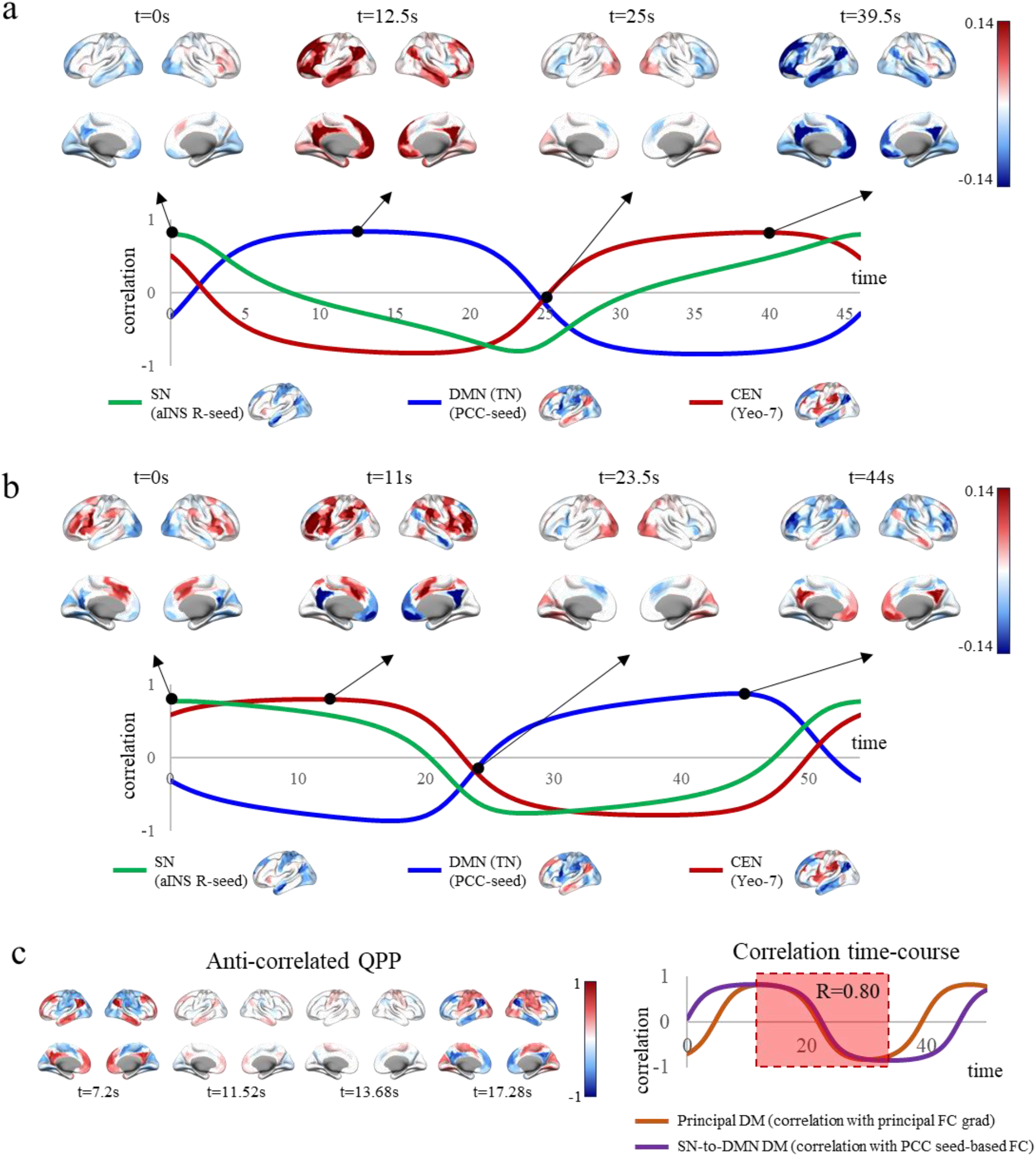
DMs 2 and 3. Reconstructed timepoints of **(a)** SN-to-DMN DM and **(b)** SN-to-CEN DM, along with the correlation time-courses between spatial patterns at each time point and the SN, DMN, and CEN. The spatial topography of SN is derived from right anterior insula (aINS R) seed-basedFC (Supplementary Figure 8a). The spatial topography of DMN is derived from PCC seed-based FC (Supplementary Figure 8b). CEN is obtained from the Yeo’s 7 network partition (Supplemenatry Figure 8c). These two DMs represent distinct transitions from SN activation: one transitions to the DMN and the other to the CEN. Both DMs exhibit anti-correlated patterns between the DMN and CEN: In the SN-to-DMN DM, DMN activation transitions to CEN; in constrast, in the SN-to-CEN DM, CEN activation transitions to DMN. **(c) Left Panel:** Spatiotemporal pattern of the anti-correlated QPP (i.e., QPP obtained with GSR). This QPP shows opposing activity between DMN (TN) and CEN (TP) regions. **Right Panel:** Correlation time-course of the principal DM and the SN-to-DMN DM. The principal DM’s time course is represented by the correlation with the principal FC gradient at each time point, while the SN-to-DMN DM represented by the correlation with the DMN (PCC seed-based FC). The combined spatiotemporal pattern of these two DMs (during both decreasing activity in transmodal regions and the DMN) highly correlates with the spatiotemporal pattern of the anti-correlated QPP (R = 0.80).

Notably, in each DM, the transition from the SN to either the DMN or the CEN was followed by subsequent activation in the previously suppressed network. For instance, in the SN-to-DMN DM, the initial activation of the SN led to increased activity in the DMN and suppression of the CEN, which was eventually followed by reactivation of the CEN. Conversely, in the SN-to-CEN DM, activation of the SN led to increased activity in the CEN and suppression of the DMN, followed by reactivation of the DMN. This was evidenced by the correlation between the spatial patterns at each reconstructed time point of these DMs and either the DMN (or TN network: PCC-seed based functional connectivity network; see Supplementary Figure 8b) or CEN (derived from Yeo’s 7-network parcellation and considered as the TP network; see Supplementary Figure 4) (Figure 3a, b). These results emphasize the inherent anticorrelation between the TN and TP networks [29, 60].

To further assess the reflection of this anti-correlation in the identified DMs, we compared them to the well-established spatiotemporal patterns of TN and TP network anti-correlation, specifically the anti-correlated QPP obtained after global signal regression (GSR) [32]. The anti-correlated QPP (see the left panel of Figure 3c) exhibited only moderate correlations with these DMs (SN-to-DMN: R = 0.54; SN-to-CEN: R = 0.41). However, when the principal DM was incorporated to reconstruct the time series, the correlations with the anti-correlated QPP increased significantly (SN-to-DMN: R = 0.80; SN-to-CEN: R = 0.59) (see the right panel of Figure 3c). This enhancement suggests that the anti-correlated QPP emerges from the combination of two distinct transitions: one from the DMN to unimodal regions (represented by the principal DM) and the other from the DMN to the CEN (represented by the SN-to-DMN DM). Thus, our findings further indicate that GSR does not eliminate the influence of the principal DM, even though reducing it significantly.

Moreover, the global signal—which reflects widespread fluctuations in brain activity—correlates more strongly with the reconstructed signals that include both the principal DM and the SN-to-DMN DM (mean R = 0.79) than with signals reconstructed using only the principal DM (mean R = 0.63; see previous section). Furthermore, the global signal showed the second-highest correlation with the global activity of the fMRI signal projected onto the SN-to-DMN DM (Supplementary Figure 5). This suggests that the principal DM remains a dominant contributor to the global signal, while the SN-to-DMN DM acts as a sub-dominant factor that further influences the global signal.

### DM4: Frontovisual-to-sensorimotor propagation in the brain

The fourth prominently engaged DM characterized by a pattern in which activity from the frontal and visual (FV) regions synchronizes and transitions into sensorimotor (SM) regions (referred to as “FV-to-SM” DM; Figure 4a), which are essential for processing sensory information and executing motor functions. We evaluated the correlation between the spatial patterns at each reconstructed time point of this DM and the second FC gradient (see the lower panel of Figure 2b for topography of the second FC gradients), which delineates functional variation along the visual-SM axis of the brain [14]. The correlation with the second FC gradient exhibited significant peaks, reaching a negative peak of −0.85 and a positive peak of 0.85. These results indicate signal propagation within this DM along the visual-SM axis, potentially reflecting the flow of information from sensory input to motor output. Regarding this, the synchronized activity between frontal regions and visual areas underscores the critical role of frontal regions in orchestrating the interaction between motor outputs and sensory inputs [61, 62].

**Figure 4.**
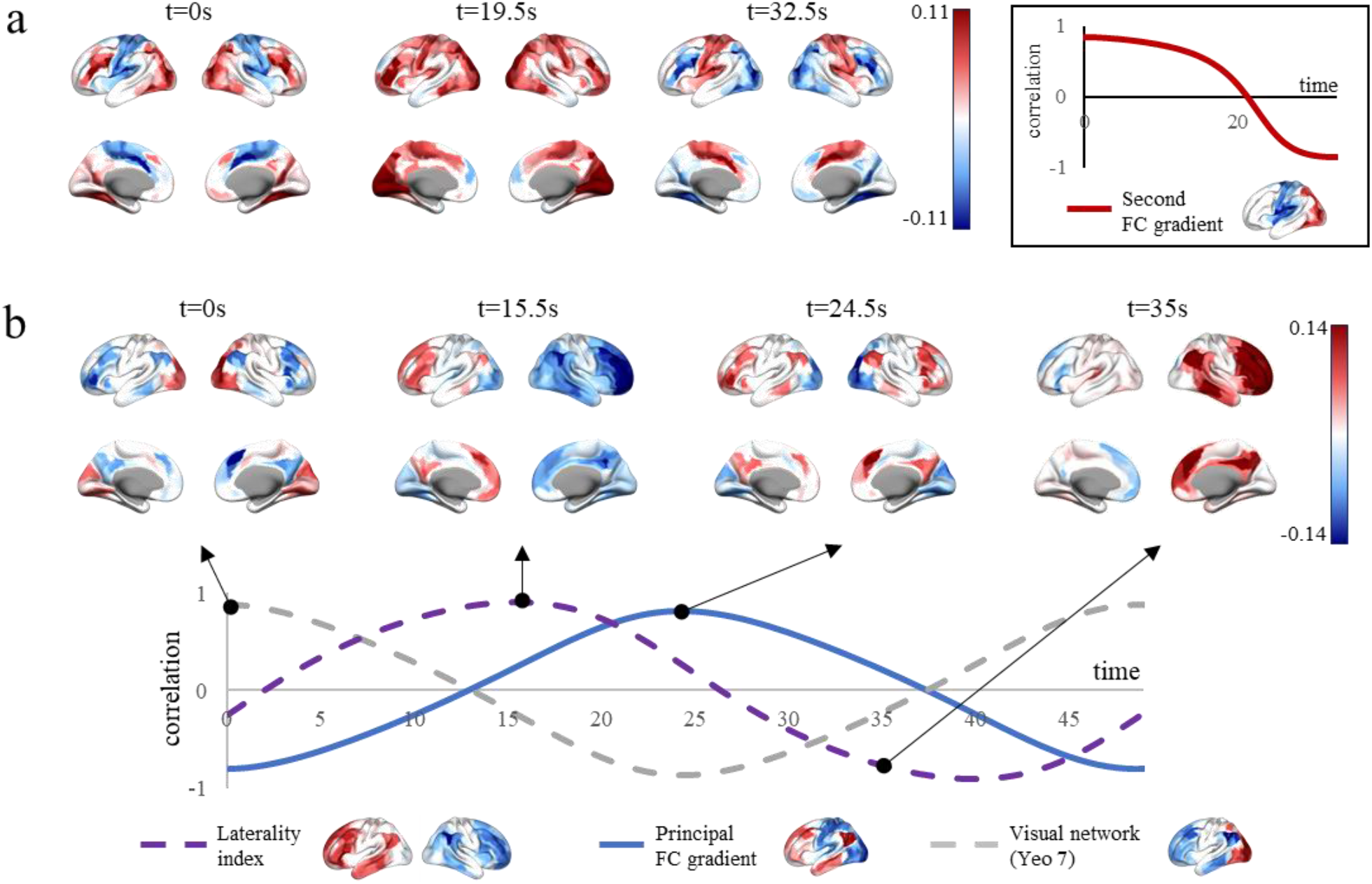
DMs 4 and 5. **(a)** Reconstructed timepoints of the FV-to-SM DM, along with the correlation time-courses between spatial patterns at each time point and the second FC gradient. This correlation indicates that signals within this DM propagate along the visual-SM axis. The synchronized activity between frontal regions and visual areas suggests that frontal regions may play a role in coordinating motor responses derived from sensory inputs. **(b)** Reconstructed time points of the bi-asym DM, along with the correlation time courses between spatial patterns at each time point and the LI, principal FC gradient, and visual network (from Yeo’s 7-network partition). In this DM, activation progresses from visual regions to left DMN regions and subsequently transitions to right DMN regions. The spatial activity patterns of the lateralization in this DM is highly correlated with the LI (R = ±0.91), indicating that this DM effectively reflects dominant hemispheric lateralization in brain dynamics. The positive correlation with the LI indicates left hemispheric dominance in activation; meanwhile, the negative correlation indicates right hemispheric dominance. The presence of this lateralized DM underscores the significance of hemispheric lateralization in brain function.

### DM5: Bilaterally asymmetric DM

Among the five DMs, the final and fifth prominently engaged DM exhibited bilateral asymmetry in its spatiotemporal pattern, highlighting the significance of hemispheric lateralization in the resting-state brain dynamics. This bilaterally asymmetric DM (bi-asym DM) initiates activation in the visual regions, subsequently progresses to the left DMN regions. The activation then transitions to transmodal regions that exhibit a robust correlation with the principal FC gradient (R = 0.81) and largely overlap with the DMN. It then shifts to the right DMN regions (Figure 4b). The correlation between the spatial pattern at each reconstructed time point and the spatial topography of laterality index (LI)—which indicates whether a ROI exhibits greater synchronization with the left or right hemisphere (see Methods)—reaches peaks of −0.91 and +0.91. This finding suggests that the bi-asym DM captures significant elements of hemispheric lateralization in resting-state brain dynamics, as quantified by the LI. Moreover, the periodic oscillations observed between the left and right hemispheres in this DM indicate an intrinsic dynamic structure that facilitates continuous interhemispheric communication.

### Characterizing brain-wide signal propagation via DM metrics: genetic influences and behavioral associations

In summary, the five DMs provide a parsimonious yet comprehensive framework for describing resting-state fMRI dynamics. Each DM is associated with specific operational functions: signal propagation along the unimodal-transmodal axis; dynamic TN/TP network switching of the SN; mapping from visual to SM regions with frontal modulation; and hemispheric lateralization in brain dynamics. Leveraging these DMs, we investigate their potential as a foundational framework for characterizing brain operations. By projecting the brain-wide fMRI signals onto the low-dimensional manifold defined by the five DMs, we effectively distill the complex spatiotemporal distributions of brain activity into their most operationally relevant components.

This dynamic projection is summarized using three distinct subject-specific metrics for each DM: (1) mode-engagement level, (2) persistence rate, and (3) progression rate. The mode-engagement level 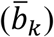 is obtained by projecting all brain-wide BOLD signals onto the DM space, reflecting the average amplitude (i.e., average engagement) of each DM during the fMRI scan. The persistence and progression rates (*r*_*k*_ and *θ*_*k*_) are derived by modeling the causal dynamics as the temporal evolution of the five DMs, characterized by subject-specific evolution coefficients 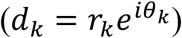. The persistence rate is given by the magnitude of the evolution coefficients, indicating how strongly a DM is sustained (i.e., how weakly decays) at the next time point. The progression rate is given by the argument (phase) of the evolution coefficients, representing the rate (i.e., speed) at which the associated neural activity patterns evolve within each mode (for mathematical details see “Group-level DMD”). Consequently, each DM yields one mode-engagement level, one persistence rate, and one progression rate, resulting in a total of 15 metrics (collectively referred to as “DM metrics”) that describe subject-specific brain-wide BOLD dynamics. The subject-averaged DM metric values are illustrated in Supplementary Figure 10a.

To assess the biological relevance and cognitive significance of these DM metrics, we examined their heritability and association with cognitive traits and behaviors. First, we investigated the genetic underpinnings of the DM metrics using twin analyses that incorporate kinship information. Heritability was estimated using the ACE model [63], which decomposes variance into additive genetic (A), common environmental (C), and unique environmental (E) components. Our results revealed significant genetic contributions to the mode-engagement levels of the five DMs (h^2^ = 0.52, 95% CI: [0.42, 0.60], p = 0.0018, n = 998), their persistence rates (h^2^ = 0.26, 95% CI: [0.17, 0.34], p = 0.0008, n = 998) and their progression rates (h^2^ = 0.36, 95% CI: [0.24, 0.48], p = 0.0002, n = 998). Further supporting these findings, we calculated the cosine distance between each pair of monozygotic (MZ) twins, dizygotic (DZ) twins, siblings, and unrelated individuals (Figure 5a). For mode-engagement levels, persistence rates, and progression rates, the five DMs exhibited greater similarity (i.e., lower cosine distance) among MZ twins compared to DZ twins (t = −3.40, Cohen’s d = −0.48, p = 4.12 × 10^−4^; t = −2.91, Cohen’s d = −0.41, p = 0.002; and t=−3.33, Cohen’s d = −0.47, p = 5.19 × 10^−4^, respectively; two-sample t-test, n = 998, one-sided). Additionally, DZ twins and siblings showed greater similarity than unrelated individuals (DZ vs. unrelated: t = −3.99, Cohen’s d = −0.41, p = 3.25 × 10^−5^; t = −2.40, Cohen’s d = −0.24, p = 0.0082; and t = −3.05, Cohen’s d = −0.31, p = 0.0012, respectively; siblings vs. unrelated: t = −5.93, Cohen’s d = −0.23, p = 1.50 × 10^−9^; t = −4.79, Cohen’s d = −0.19, p = 8.25 × 10^−7^; and t = −6.88, Cohen’s d = −0.27, p = 2.92× 10^−12^, respectively; two-sample t-test, n = 998, one-sided). No significant differences were observed between DZ twins and siblings (p = 0.054, p = 0.30 and p = 0.36, respectively; two-sample t-test, n = 998, one-sided).

**Figure 5.**
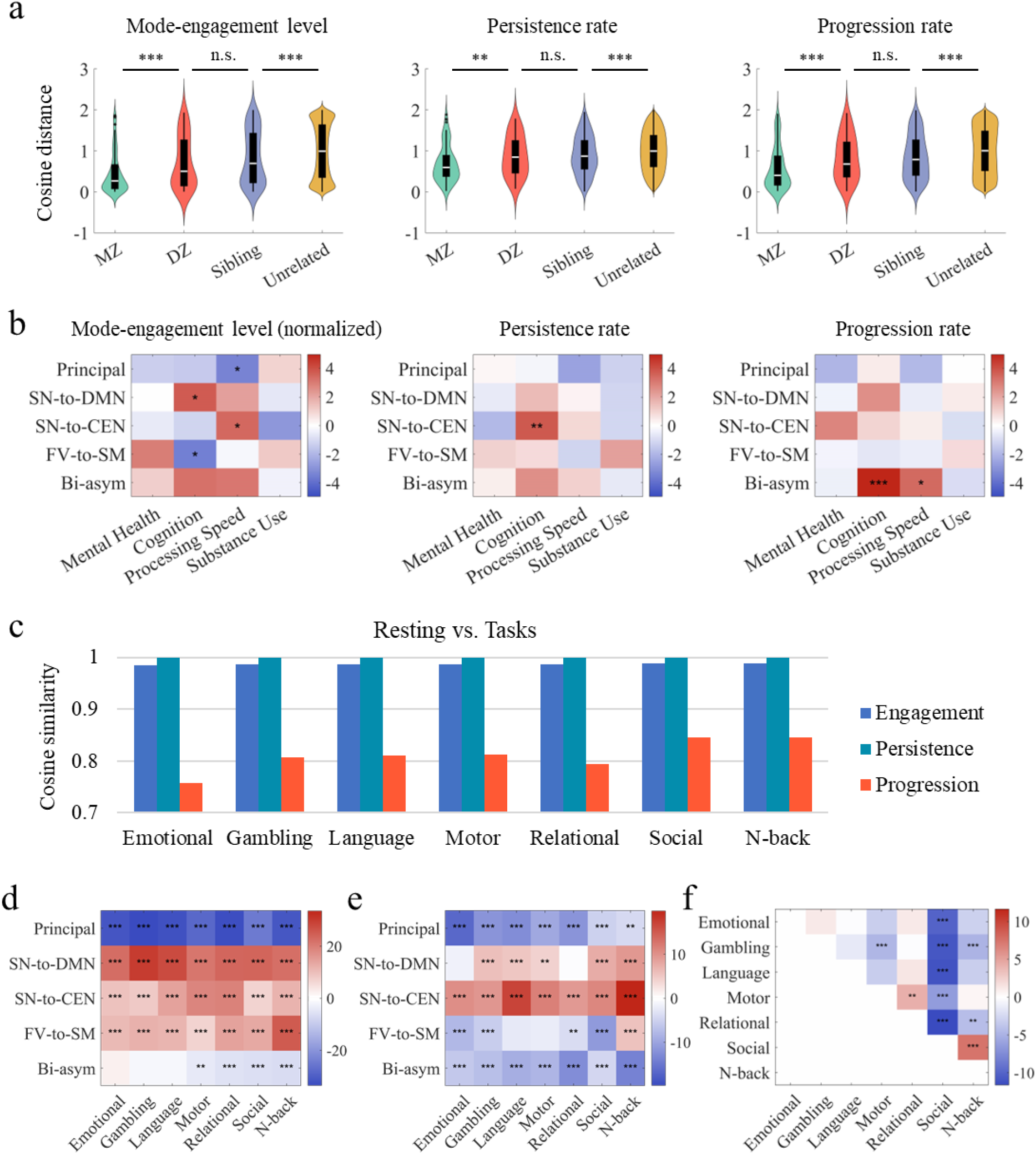
Influences of genetics, behavior, and task. **(a)** Cosine distances between DM metrics for each pair of MZ twins, DZ twins, siblings, and unrelated individuals (n = 998). Distances were calculated separately for mode-engagement levels, persistence rates, and progression rates due to differences in value ranges and distributions. The plotted distances represent residuals after regressing out confounding variables (e.g., age, gender, years of education, and others; see Methods). The statistical significance of the differences between groups was assessed using a one-sided two-sample t-test. **(b)** T-value heatmap illustrating the correlation between four behavioral latent factors—mental health, cognition, processing speed, and substance use—and the DM metrics (n = 1,086). P-values were FWE-corrected (family-wise error corrected) for all 60 tests (combinations of three DM metrics, five DMs, and four behavioral factors) via permutation methods (see Methods). Mode-engagement levels were normalized so that the sum of the five values for each subject equals 1, facilitating the assessment of each DM’s engagement ratio (referred to as normalized mode-engagement level). See Supplementary Tables 1-6 for the detailed statistical results. **(c)** Similarity of DM metrics between resting and task states (n = 994). Similarities were evaluated separately for mode-engagement levels, persistence rates, and progression rates due to differences in value ranges and distributions. All three metric types exhibited high similarity between resting and task states (CS > 0.7). **(d–f)** T-value heatmaps comparing DM metrics between resting and task states: **(d)** Normalized mode-engagement rates in resting versus task states, **(e)** Progression rates in resting versus task states, and **(f)** Normalized mode-engagement rates of the principal DM across all pairs of seven tasks (n = 994). In comparisons between resting and task states, positive t-values (red) indicate higher DM metrics during tasks (labeled on the column) compared to rest, while negative t-values (blue) indicate lower metrics during tasks. In comparisons among task states, positive t-values (red) signify that the DM metric for the task labeled on the row is significantly greater than that for the task labeled on the column, and negative t-values (blue) indicate the opposite. P-values for resting vs. task comparisons were Bonferroni-corrected separately for the three DM metric types. For task state comparisons, p-values were Bonferroni-corrected across all five DMs and all possible pairs of the seven tasks (resulting in 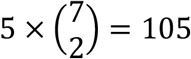 comparisons), while still correcting separately for the three types of DM metrics. See Supplementary Figures 10–13 for additional comparison results. Asterisks indicate statistical significance (*p < 0.05, **p < 0.01, ***p < 0.001).

Next, to explore the cognitive significance of the DM metrics, we examined their correlations with participants’ cognitive traits and behaviors. We utilized previously established latent factors derived from behavioral items in the HCP dataset [64]: (1) mental health, (2) cognition, (3) processing speed, and (4) substance use. Correlation analyses were conducted between these four behavioral factors and the DM metrics, controlling for age, sex, handedness, and the cube root of intracranial volume (ICV) and total gray-matter volume (TGMV) as confounding variables (Figure 5b). We found that the cognition factor, reflecting intellectual abilities across various domains, was positively correlated with the mode-engagement level of the SN-to-DMN DM (t = 3.69, p = 0.0148, FWE-corrected, n = 1,086, two-sided) and negatively correlated with the mode-engagement level of the FV-to-SM DM (t = −3.51, p = 0.0285, FWE-corrected, n = 1,086, two-sided). Similarly, the processing speed factor, reflecting efficiency of cognitive processing, was positively correlated with the mode-engagement level of the SN-to-CEN DM (t = 3.42, p = 0.0382, FWE-corrected, n = 1,086, two-sided) and negatively correlated with the mode-engagement level of the principal DM (t = −3.39, p = 0.0434, FWE-corrected, n = 1,086, two-sided). These results indicate that a higher presence of the SN-to-DMN and SN-to-CEN DMs in BOLD dynamics is associated with better performance of many cognitive tasks, emphasizing the significance of these DMs in cognitive functioning.

The persistence and progression rates also showed correlations with the cognition and processing speed factors. Specifically, the persistence rate of the SN-to-CEN DM was positively correlated with the cognition factor (t = 3.77, p < 0.0117, FWE-corrected, n = 1,086, two-sided), further indicating the significance of this DM in task performance. The progression rate of the bi-asym DM was positively correlated with both the cognition factor (t = 4.94, p < 1 × 10^−4^, FWE-corrected, n = 1,086, two-sided) and processing speed (t = 3.39, p = 0.0432, FWE-corrected, n = 1,086, two-sided), supporting the notion that lateralized dynamics are related to cognitive abilities. These results underscore that dynamic features—such as the degree to which the involvement of DMs is sustained (as indicated by the persistence rate) and the speed at which they evolve (as indicated by the progression rate)—are also related to cognitive abilities. Detailed statistical results for the behavioral analyses are provided in Supplementary Tables 1-6. Overall, the dynamical characteristics represented by the DM metrics reflect important latent factors associated with general cognitive performance.

### Individual signatures of DMs are largely preserved during tasks

We further investigated whether the overall signatures of DMs remain consistent during task performance. To test this, we analyzed task-state fMRI data from the HCP dataset, which includes seven distinct tasks: emotional processing, gambling, language comprehension, motor execution, relational reasoning, social cognition, and working memory (N-back) tasks [65]. Each task engages different basic cognitive aspects, providing a comprehensive assessment of brain dynamics under varying cognitive demands.

Our analysis confirmed that all five DMs were significantly present during every task (t > 6.8, p < 1 × 10^−10^; see Methods), underscoring the robust presence of these core dynamic structures across different task conditions. Furthermore, we observed that the relative patterns of both magnitude and phase for the five DMs closely matched those seen during the resting state. Specifically, the cosine similarity (CS) values exceeded 0.98 for mode-engagement levels, greater than 0.999 for persistence rates, and over 0.75 for progression rates when comparing task states to rest (Figure 5c). These high CS values demonstrate a remarkable similarity between resting-state and task-state DMs, supporting the notion that the brain’s intrinsic dynamic architecture is preserved during task execution [66]. Therefore, these results indicate that the overall signatures of DMs are stable features of brain activity that persist even as cognitive demands change.

Moreover, we found that the CS values for DM metrics within the same subject across different tasks were all significantly higher than the CS values between different subjects performing the same task. For the mode-engagement levels, the average CS between different tasks for the same subject was 0.991, which was significantly higher (i.e., more similar) than the average CS between different subjects performing the same task (average CS = 0.989; two-sample t-test, 95% CI for difference = [0.0018, 0.0021], t = 31.55, p = 2.01 × 10^−218^, Cohen’s d = 0.22, n = 994, two-sided). Similarly, the persistence and progression rates for the same subject between different tasks (persistence rate: average CS = 0.999716; progression rate: average CS = 0.747) were significantly more similar than those for the same task between different subjects (persistence rate: average CS = 0.999684, two-sample t-test, 95% CI for difference = [2.79 × 10^−05^, 3.60 × 10^−05^], t = 15.39, p = 1.95 × 10^−53^, Cohen’s d = 0.11, n = 994, two-sided; progression rate: average CS =0.728, two-sample t-test, 95% CI for difference = [0.015, 0.021], t = 11.02, p = 2.91 × 10^−28^, Cohen’s d = 0.08, n = 994, two-sided). In sum, individual differences in DMs are consistent across different tasks, highlighting the stability of individual DM signatures.

While the overall signatures of DMs are preserved during tasks, we also observed task-dependent changes in the magnitude and phase of specific DMs. Specifically, the mode-engagement level of the principal DM was significantly reduced during task execution compared to the resting state; in contrast, the SN-to-DMN, SN-to-CEN, and FV-to-SM DMs exhibited increased engagement (Figure 5d), highlighting their enhanced involvement in task-related processes. The progression rates of the SN-to-DMN and SN-to-CEN DMs increased during all seven tasks compared to rest, while other DMs mostly decreased (Figure 5e), implicating changes in the anti-correlation between TP and TN networks during tasks.

Furthermore, we detected significant variations in DMs between different tasks, suggesting that specific cognitive demands can modulate brain dynamics. For instance, the principal DM was significantly more engaged during social tasks, which involve interpreting social interactions (Figure 5f). This increased engagement may reflect the necessity for integrating information across widespread brain regions to comprehend complex social cues. Conversely, the principal DM exhibited greater persistence during N-back tasks, which require heightened working memory and sustained attention (Supplementary Figure 12). This increased persistence may indicate the need for maintaining stable neural representations over time to support working memory processes. Additional differences are provided in Supplementary Figures 11–13. These findings indicate that while the core DMs are maintained, their relative contributions vary depending on the nature of the task, highlighting the flexibility of the brain’s dynamic architecture in adapting to specific cognitive demands.

## Discussion

Here, we identified five large-scale signal propagation modes (i.e., DMs) that establish a parsimonious schema of the causal dynamics in the brain. These modes achieve a balance between richness and simplicity in explaining whole-brain activity: their superposition effectively predicts future trajectories of BOLD signals. The discovery of these five DMs significantly advances our understanding of how the brain operates by resolving macroscale dynamics into fundamental units that evolve over time. Each DM aligns with dynamic reconfigurations of established large-scale networks, integrating temporally distinct yet functionally connected network configurations and elucidating mechanisms governing state-transition trajectories. We further showed that this framework unifies various canonical features of brain dynamics: they can emerge directly from interactions among these five DMs. By projecting brain-wide signals onto these five DMs, we derived interpretable metrics quantifying mode-specific engagement, persistence, and propagation speed. Individual differences in these metrics were associated with domain-general cognitive abilities, were influenced by genetic factors, and remained consistent across various tasks, underscoring their relevance to behaviorally meaningful neural dynamics.

The principal DM (DM1) captures the dominant signal propagation within the brain, reflecting the intrinsic flow of information between unimodal sensory regions and transmodal association areas [11–13]. This observation likely signifies multisensory integration in transmodal regions and top-down modulation to unimodal areas. Furthermore, this finding is consistent with earlier descriptions of transitions between extrinsic and intrinsic systems as two temporal “metastates” [2] and aligns with the first principal component identified in prior applications of complex PCA to fMRI datasets [67]. These converging lines of evidence highlight the principal DM’s significant role in brain dynamics, as revealed by various spatiotemporal pattern detection methodologies. The robust association between DM1 and key spatiotemporal structures—such as the principal functional gradient encompassing the cortex, subcortical areas, and cerebellum [14, 49]; wave propagation patterns [11–13]; transitions between metastates [2]; global signal [50]; lag structure [51]; and QPP [32]—further highlights the dominance of this signal propagation. Additionally, these associations suggest that the propagation characterized by the principal DM serves as a common source for these diverse spatiotemporal phenomena. The persistence of this dominant mode during task performance indicates its potential involvement in goal-directed and executive cognitive functions.

A notable finding of our study is the identification of propagation modes reflecting the dynamic interplay among the three major large-scale networks namely, the DMN, CEN, and SN—captured in DM2 and DM3 (SN-to-DMN and SN-to-CEN DMs). The activation of the SN can lead to engagement of either the DMN or the CEN, potentially modulating transitions between internally focused and externally oriented cognitive states. This finding underscores the SN’s role as a pivotal switching hub between the DMN and CEN, as proposed in the well-established triple-network model [35, 37]. Furthermore, the involvement of the DMN and CEN was followed by activation of the opposing network, indicating the well-known anticorrelation between TN and TP networks (i.e., DMN and CEN) [29, 68]. The relative dominance of these DMs positively correlated with domain-general cognitive abilities (Figure 5b), aligning with previous reports associating strong anticorrelation of TN/TP networks with flexible task-related resource allocation [69, 70]. This suggests that signal propagation between these networks is crucial for task-related processes, and that modulation by the SN plays a significant role.

The FV-to-SM DM (DM4) represents the synchronization and propagation of activity from frontal and visual regions to SM areas. This associated with the second cortical FC gradient, which varies along the visual-SM axis [14], thereby highlighting the integration of sensory inputs with motor outputs essential for coordinated action. The observed synchronization between frontal and visual regions suggests that modulation within frontal regions plays a critical role in this integration.

Finally, the bi-asym DM (DM5) reveals bilaterally asymmetric dynamics, emphasizing the significance of hemispheric lateralization in cognitive processing and inter-hemispheric communication. The propagation within this DM strongly reflects the LI, demonstrating that this single method (i.e., DMD) can also capture the prominent lateralized spatiotemporal patterns previously identified through other sophisticated methodologies [52]. Signal propagation within this DM oscillates between the left and right hemispheres, indicating a pattern of interhemispheric communication. The observed propagation speed (as indicated by the progression rate) suggests facilitation of interhemispheric communication, which is considered a crucial factor for high-level cognitive functions (e.g., working memory) and processing speed [71–73]. Consistent with this, the propagation speed of this DM was associated with higher cognitive performance and processing speed (Figure 5b), thereby suggesting that enhanced interhemispheric communication is linked to superior cognitive abilities. It is noteworthy that only this DM is unique in exhibiting a behavioral association with propagation speed. Furthermore, the involvement of the visual network and the DMN during the interhemispheric transitions may signify their critical roles in this communication process, highlighting the necessity for further investigation in future studies.

Notably, the overall signatures of the DMs (represented by DM metrics) are genetically influenced and largely preserved across different task conditions. The DM metrics are more similar within subjects across tasks than between subjects on the same task, underscoring stable aspects of the brain’s functional topology [66]. However, we also observed task-dependent modulations in DMs, characterized by increased involvement and facilitation in the SN-to-DMN and SN-to-CEN DMs, alongside a relative decrease in the principal and FV-to-SM DMs. This aligns with the operational interpretation of the SN-to-DMN and SN-to-CEN DMs as being involved in task-related processing, potentially capturing the task-modulation of anticorrelation of TN/TP networks [31, 74]. Overall, these findings enhance our understanding of how the brain maintains core functional dynamics while adapting to varying cognitive demands.

Among the five DMs, four (the principal DM, SN-to-DMN DM, SN-to-CEN DM, and bi-asym DM) are associated with the DMN. This widespread association underscores the DMN’s pivotal role as a central functional hub in the brain, responsible for integrating and coordinating diverse types of information across various large-scale networks [25, 75–78]. However, we noted minor differences in DMN activation among these DMs. For instance, the ventromedial prefrontal cortex (VMPFC) is rather deactivated (activation value: –0.0547 at area 10v in the Glasser atlas) during DMN-like activation in the principal DM (Figure 2a), but it is activated (activation value: > 0.14 at area 10v) during DMN-like activation in the SN-to-DMN and SN-to-CEN DMs (Figure 3a, b). Further differences were observed in the ventrolateral PFC and temporal regions during DMN-like activation in these DMs. These variations may reflect operational distinctions among DMN nodes: for example, the VMPFC might be more relate to task-related processing [79, 80]. This could explain the division of subnetworks within the DMN [81, 82], warranting further exploration in future studies.

Our framework also offers an interpretation for global signals in the brain, a topic of extensive discussion in the neuroimaging field [50, 83–86]. Our observations indicate that the major contributor to the global signal is the propagation along the unimodal-transmodal axis, as indicated by the principal DM, but it is also significantly influenced by the anticorrelation between task-negative and task-positive networks in the SN-to-DMN DM. Consequently, GSR significantly reduces the predominant propagation along the unimodal-transmodal axis from the original fMRI time series, making subsequent analyses more sensitive to other signal propagations in the brain [84]. However, our findings suggest that GSR does not entirely remove the influence of the principal DM, as evidenced by the significant explanatory ability of the principal DM in anticorrelated QPPs even after GSR. Therefore, our observations offer new insights into the ongoing debates surrounding the interpretation of global signals and the use of GSR in neuroimaging.

It is important to note that the five DMs we focused on are not exhaustive patterns of signal propagation in the brain. Nonetheless, we believe that our conceptual framework remains robust: whole-brain dynamics can be effectively captured through the superposition of a meticulously selected set of signal propagation modes. Although additional DMs identified in the group-level DMD do not enhance the predictive ability for future BOLD signals, these modes may be potential candidates for nuisance aspects of brain dynamics, which should be addressed in future studies. Moreover, one limitation of projecting onto DMs is that the estimation of DM metrics depends on the presence of other DMs during projection due to the non-orthogonality of DM vectors. Therefore, characterizing the residual projections associated with additional DMs is essential for accurately capturing an individual’s brain dynamics within this DM framework.

In conclusion, we resolve the critical challenge of capturing rich temporal sequences in a parsimonious modeling framework of brain dynamics by identifying fundamental propagation modes (DMs). Our work lays a groundwork for moving beyond static “brain states” by formalizing dynamics as interdependent wave-like motifs [9], thereby reconciling the brain’s stable temporal topology with its capacity for adaptive network reconfiguration. Through a superposition framework that integrates various spatiotemporal features, we demonstrate how cortical activity arises from a superposition of waves traversing a continuous neural medium—a direct embodiment of neural field theory’s core principles [87–89]. The DM-based approach offers a streamlined approach for characterizing individual neural fingerprints, suggesting its translational potential for representing alterations in patients’ brain dynamics.

## Methods

### HCP dataset and preprocessing

We employed both resting-state and 7-task fMRI data from the HCP S1200 release [90]. For each participant, resting-state fMRI scans were conducted over the course of two days. Each day included two separate 15-minute resting-state sessions (with eyes open and fixed) utilizing opposite phase-encoding directions (L/R: left-to-right, R/L: right-to-left), resulting in a total of four resting-state scans per individual. Following the resting-state scans each day, participants completed 30 minutes of task fMRI data collection. The seven tasks—emotional, gambling, language, motor, relational reasoning, social, and working memory (N-back)—distributed across the two sessions, with each task administered twice using opposing phase-encoding directions (L/R and R/L). Detailed descriptions of the resting-state and task fMRI data collection protocols are available in prior publications [65, 91].

For the resting-state analysis, we included participants who completed at least one day of resting-state scanning, which comprised two resting-state scans with opposing phase-encoding directions. This selection resulted in a sample of 1,086 subjects (female = 590, mean age = 28.78, std = 3.70). The number of subjects for each task fMRI varied; therefore, we included all participants who completed two opposing phase-encoding sessions for each specific task. The sample sizes for assessing the significance of DMs during tasks were as follows: emotional (n = 1,043, female = 560, mean age = 28.77, std = 3.70), gambling (n = 1,078, female = 582, mean age = 28.77, std = 3.70), language (n = 1,045, female = 559, mean age = 28.75, std = 3.69), motor (n = 1,079, female = 584, mean age = 28.76, std = 3.69), relational reasoning (n = 1,038, female = 555, mean age = 28.75, std = 3.70), social (n = 1,048, female = 562, mean age = 28.76, std = 3.69), and working memory (n = 1,079, female = 586, mean age = 28.78, std = 3.69). Among them, 994 participants (female = 530, mean age = 28.71, std = 3.71) completed at least one day of resting-state scanning and all seven tasks, which were used to compare the similarities and differences of DMs between the resting state and the seven tasks.

All imaging data were collected at Washington University in St. Louis using a customized Siemens 3T Skyra scanner, which utilized a multiband sequence for rapid image acquisition. Structural images were obtained at 0.7-mm isotropic resolution. Resting-state fMRI data were collected with 2-mm isotropic spatial resolution and a repetition time (TR) of 0.72 seconds. Further details regarding data acquisition and preprocessing pipelines are described elsewhere [90, 91]. Informed consent was obtained from all participants, and all procedures adhered to relevant ethical guidelines and regulations.

For the resting-state data, we utilized surface-based CIFTI fMRI scans that were registered using MSMAll and had been preprocessed with the HCP’s ICA-based artifact removal process (ICA-FIX) to minimize the effects of spatially structured noise [91]. For the task-based data, we utilized MSMAll-registered fMRI scans that were minimally preprocessed.

Both resting-state and task-state fMRI scans were spatially parcellated using a comprehensive brain-wide parcellation scheme. This scheme combined the Glasser cortical parcellation [53] with the Cole-Anticevic Brain-wide network partition for subcortical and cerebellar regions [54]. This approach resulted in a set of ROIs covering both cortical and subcortical areas, specifically yielding 360 cortical parcels and 358 subcortical and cerebellar parcels. However, we identified two ROIs from the Cole-Anticevic Brain-wide Network Partition that exhibited negligible variance (variance < 0.001), indicating a lack of data in these regions. These ROIs were key value 1040 labeled “Visual1-14_R-Caudate” and key value 7290 labeled “Frontoparietal-29_R-Cerebellum.” Consequently, we excluded these two ROIs from further analysis. In summary, our final parcellation scheme consisted of 360 cortical ROIs and 356 subcortical and cerebellar ROIs, totaling 716 regions that were used for subsequent analyses.

Temporal preprocessing involved applying a zero-phase Butterworth band-pass filter with a frequency range of 0.01–0.1 Hz to the data to eliminate low frequency signal drifts and physiological noise [92, 93]. For the task-state fMRI scans, we additionally regressed out 24 motion parameters, linear and quadratic trends, and task-related activations before band-pass filtering to control for head motion artifacts and the mere effects of brain-wide task-related coactivation. The task-related activations were modeled by convolving boxcar regressors—constructed from task onset and duration information—with a hemodynamic response function. This regression was performed using the ordinary least squares method. No temporal normalization (e.g., z-scoring or variance normalization) was applied to either the resting-state or task-state datasets to preserve the original amplitude of the BOLD signals.

### Group-level DMD

We first estimated the group-level causal connectivity matrix *A* ∈ ℝ^*N*×*N*^, which best describes the causal relationships between temporally adjacent BOLD activations across all subjects:

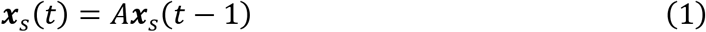

where ***x***_*s*_(*t*) ∈ ℝ^*N*^ represents the BOLD activation vector at time t for the *s*-th subject, and *N* denotes the number of ROIs.

Our objective was to find the matrix *A* that minimizes the cumulative prediction error across all subjects and time points:

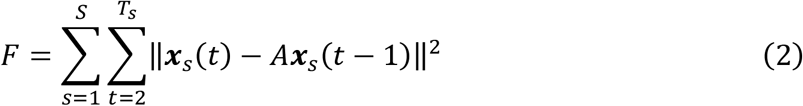

where *S* is the total number of subjects, *T*_*s*_ is the number of time points for subject *s*, and ‖·‖ denotes the Euclidean norm.

To solve this optimization problem efficiently, we constructed two data matrices by concatenating the BOLD activation vectors across all subjects and time points: 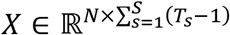 constructed by stacking ***x***_*s*_(*t* − 1) for all subject *s* ∈ {1,2, ⋯, *S*} and time points *t* ∈ {2,3, ⋯, *T*} and 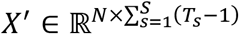 constructed by stacking ***x***_*s*_(*t*). Specifically, the two matrices are like the followings:

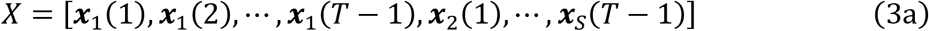

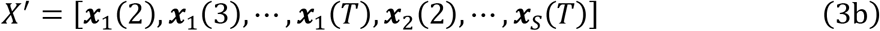

The optimal matrix *A* that minimizes *F* can then be obtained using the least-squares solution [44, 94]:

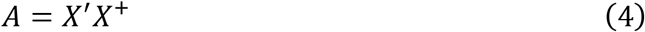

where *X*^+^ is the Moore-Penrose pseudoinverse of *X*.

However, measurement noise can introduce significant systematic bias in the DMD results because the above approach only minimizes the errors in *X*^′^, implicitly assuming that there is no measurement error in *X*. To mitigate this bias, we employed the forward-backward DMD algorithm, which provides a simple yet effective method to reduce noise-induced bias [55]. The forward-backward DMD approach operates on the premise that the effect of measurement noise on DMD is stable and systematic. In addition to estimating the forward dynamics ***x***_*s*_(*t*) = *A****x***_*s*_(*t* − 1), we also approximate the inverse dynamics:

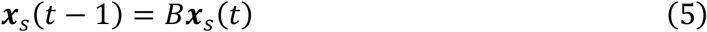

where *B* ∈ ℝ^*N*×*N*^ is another matrix to be estimated.

By computing both *A* and *B*, we can calculate a new, less-biased estimate of the system dynamics:

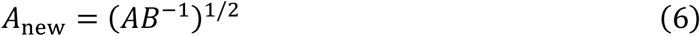

where (·)^1/2^ denotes the matrix square root. This method leverages the systematic but opposite biases in A and B to achieve a more accurate estimation of the true system’s DMs. To compute *B*, we utilized the data matrices in reverse order and solved:

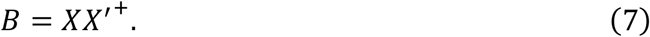

Finally, we calculated the improved estimate *A*_new_ using equation (6) and performed an eigenvalue decomposition to obtain the group-level DMs (***ϕ***_*k*_, complex eigenvectors) and their evolution coefficients (*λ*_*k*_, corresponding complex eigenvalues):

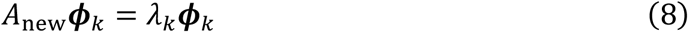

where *k* is eigenvalue index. Since *A*_new_ is a real value matrix, its complex eigenvalues and eigenvectors occur in conjugate pairs. This property ensures that when reconstructing the BOLD signals, the imaginary parts cancel out, resulting in real-valued activations. Each pair of conjugate eigenvectors corresponds to one DM, capturing oscillatory patterns in the data [57]. Prior to applying DMD, we down-sampled the fMRI data from the original TR of 0.72 seconds to a TR of 1.5 seconds using cubic spline interpolation (implemented in MATLAB) to reduce data complexity while enhancing the detection of meaningful dynamic differences due to the inherently slow hemodynamic responses.

The complex eigenvalues *λ*_*k*_, referred to as “group-level (temporal) evolution coefficients” of the *k*-th DM, can be expressed in polar form using two real numbers: magnitude *r* and phase *θ*, such that 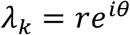.Due to the properties of eigenvectors, if the initial BOLD activation is ***x***_*s*_(0) = ***ϕ***_*k*_, then after 1TR, ***x***_*s*_(1) = *A****ϕ***_*k*_ = *λ*_*k*_***ϕ***_*k*_ = *re*^*iθ*^***ϕ***_*k*_; after 2TR, ***x***_*s*_(2) = *A****x***_*s*_(1) = *A*(*λ*_*k*_***ϕ***_*k*_) = *λ*_*k*_^2^***ϕ***_*k*_ = *r*^2^*e*^2*iθ*^***ϕ***_*k*_; and so on.

In this context, the magnitude of the evolution coefficient *r* indicates the proportion of the signal that persists after one TR, reflecting the stability or sustained influence of the DM. We will refer to this as “*persistence rate*.” A higher persistence rate indicates that the DM maintains its influence over time, suggesting more stable or enduring neural activity within that mode. Meanwhile, the phase *θ* determines the rate at which the DM evolves. Specifically, the period of oscillation is given by 2*π*/*θ* TRs, which quantifies the speed of signal propagation. We will refer to the phase *θ* as “*temporal progression rate*” or “*progression rate*.” A higher temporal progression rate signifies faster oscillatory dynamics, indicating quicker changes in the neural activity patterns associated with that mode.

### Subject-level estimation of DM metrics

Building upon the understanding that the fundamental organizational principles and large-scale network architectures of the human brain are remarkably consistent across individuals [95], we modeled the subject-level fMRI dynamics using the same set of large-scale signal propagation modes [94]—i.e., DMs derived from group-level analysis—across all individuals, while allowing for variations in the propagation strength and speed unique to each subject.

To this end, we decomposed each subject’s causal BOLD signal dynamics into the temporal evolution of the DMs. Specifically, for each subject *s* and given *K* DMs, the BOLD signal at time *t* − 1, ***x***_*s*_(*t* − 1), can be decomposed as:

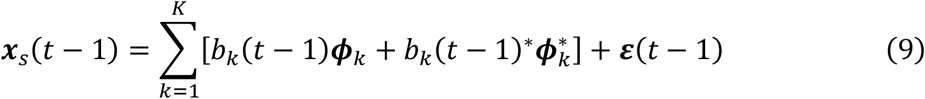

where the coefficient *b*_*k*_(*t* − 1), referred to as “projection coefficient”, of group-level DM vector ***ϕ***_*k*_ is estimated by projecting the BOLD activation vector ***x***_*s*_(*t* − 1) onto the DM vector space, reflecting engagement level of the *k*-th DM’s activation at time *t* − 1. *b*_*k*_(*t* − 1)^∗^ and 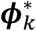 are complex conjugate of *b*_*k*_(*t* − 1) and ***ϕ***_*k*_. The residual ε(*t* − 1) is the orthogonal component remaining after this projection of ***x***_*s*_(*t* − 1).

Assuming that the global structure of signal propagation for each DM is preserved, the subsequent BOLD signal can be modeled as:

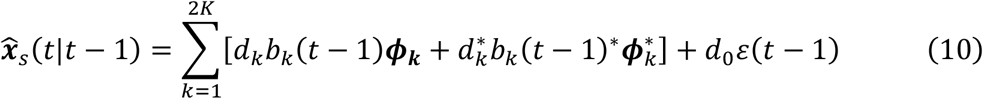

where *d*_*k*_ are subject-specific evolution coefficients representing the persistence and progression rates for each mode in that subject’s brain (see previous section), and *d*_0_ is an additional subject-specific (autocorrelation) coefficient accounting for autocorrelation in the residual 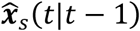 is the predicted BOLD signal at time t based on information up to time *t* − 1. Essentially, we modeled the subject-level fMRI dynamics using the same large-scale signal propagation modes across individuals, allowing for variations in propagation strength and speed. Our objective is therefore to determine the evolution coefficients *d*_*k*_ and 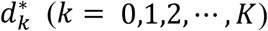 that best approximate the predicted signal 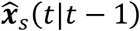 to the actual observed signal ***x***_*s*_(*t*).

To solve this problem, we reorganized it using matrix notation. Let Φ ∈ ℂ^*N*×*M*^ denote the complex matrix whose columns are the eigenvectors corresponding to the selected DMs, where *N* is the number of ROIs and *M* = 2*K* accounts for complex conjugate pairs of eigenvectors (since the eigenvalues and eigenvectors of real matrices appear in complex conjugate pairs).

For each subject *s*, we approximated their fMRI causal dynamics as a combination of the selected DMs modulated by subject-specific evolution coefficients. Specifically, we modeled the dynamics as:

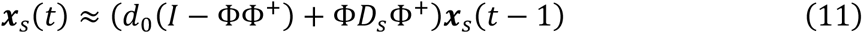

where ***x***_*s*_(*t*) ∈ ℝ^*N*^ represents the BOLD activation vector at time t for the *s*-th subject, *I* is the identity matrix of size *N*-by-*N*, 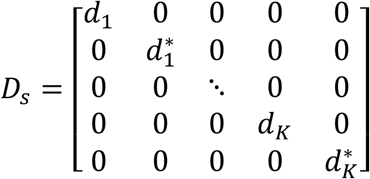 is a diagonal matrix containing the subject-specific evolution coefficients *d*_*k*_, and their complex conjugates. *d*_0_ represents the autocorrelation coefficient of the orthogonal complement of the column space of Φ. In this formulation, ΦΦ^+^ is the projection matrix onto the subspace spanned by the selected DMs, and (*I* − ΦΦ^+^) projects onto the orthogonal complement of that subspace. Thus, the term *d*_0_(*I* − ΦΦ^+^)***x***_*s*_(*t* − 1) accounts for the residual dynamics not captured by the selected DMs.

Our objective was to estimate the evolution coefficients *d*_*k*_ for *k* = 0,1, ⋯, *K* that minimize the cumulative prediction error across all time points for each subject:

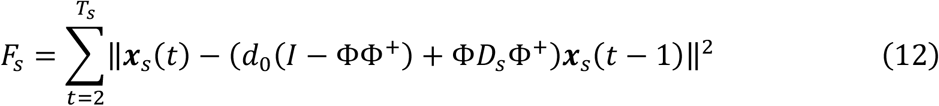

where *T*_*s*_ is the total number of time points for subject *s*.

To minimize *F*_*s*_, we formulated the problem as solving a system of linear equations. This was achieved by expanding *F*_*s*_ and setting the partial derivatives (see Supplementary Result 2) with respect to each *d*_*k*_ to zero, resulting in the following linear system:

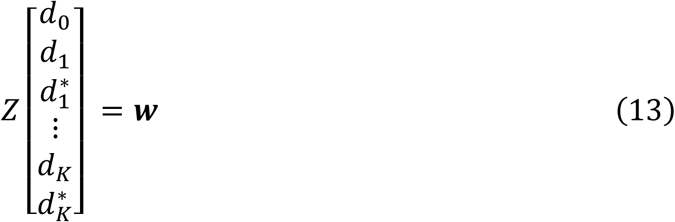

where *Z* ∈ ℂ^(2*K*+1)×(2*K*+1)^ is a square matrix, and ***w*** ∈ ℂ^(2*K*+1)^ is the right-hand side vector.

The elements of *Z* and ***w*** are defined based on the data ***x***_*s*_(*t*) and the projections onto the subspace spanned by Φ and its orthogonal complement.

For *Z* (*i, j* ∈ [1,2, ⋯, 2*K*]):

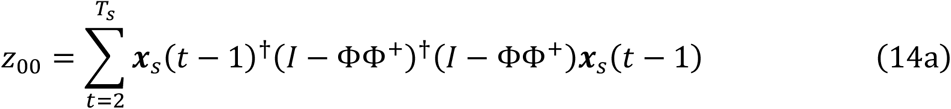

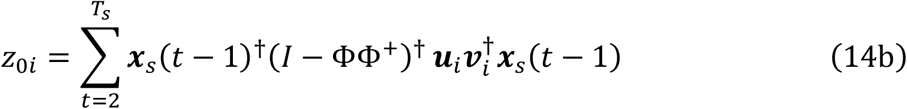

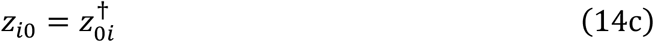

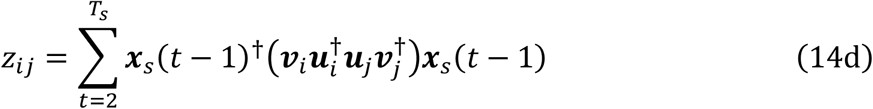

For ***w*** (*i* ∈ [1,2, ⋯, 2*K*]):

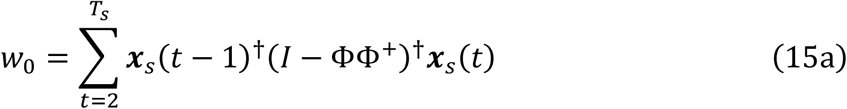

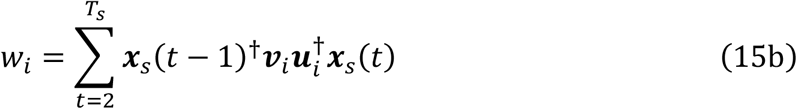

where ***u***_*i*_ is *i-*th column vector of Φ, and ***v***_*i*_ is *i-*th column vector of (Φ^+^)^†^, where † denotes conjugate transpose. By solving this linear system for each subject, we obtained the subject-specific coefficients:

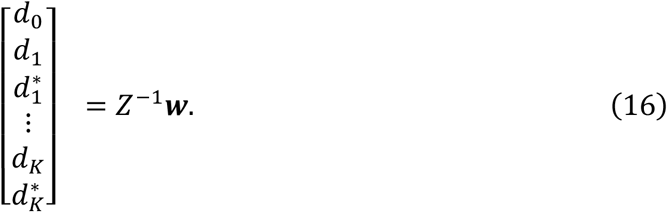

The detailed derivation of the linear system and the definitions of *Z* and ***w*** are provided in Supplementary Result 2.

Subsequently, we can derive the subject-specific persistence rate (*r*_*k*_) and progression rate (*θ*_*k*_) can be achieved from the evolution parameters, expressed as 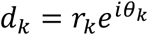 for *k* = 1,2, ⋯, *K*. It is important to note that while the persistence rates from *d*_*k*_ and its complex conjugate 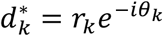 are same as *r*_*k*_, the progression rates have opposite signs (*θ*_*k*_ and −*θ*_*k*_, respectively). Therefore, we exclusively use *d*_*k*_ (i.e., not 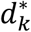) for estimating the DM metrics.

Additionally, we estimated the subject-specific *mode-engagement level*, 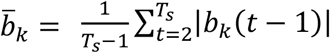, which indicates how dominantly the *k*-th DM is engaged during the fMRI scan. The projection coefficient, *b*_*k*_(*t* − 1), can be obtained by applying Φ^+^ to ***x***_*s*_(*t* − 1):

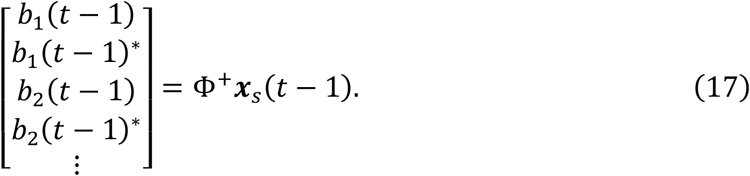

For the resting-state data, we utilized the down-sampled fMRI scans with a TR of 1.5 seconds, as described in the previous section. In contrast, we did not down-sample the task-state data due to the limited number of time points available in these datasets. Therefore, when comparing the DM coefficients between the resting-state and task-state data, we adjusted the resting-state DM coefficients to ensure temporal consistency between the two datasets. Specifically, we scaled the resting-state DM coefficients by raising them to the power of 0.72/1.5, accounting for the difference in TRs (1.5 seconds for resting-state and 0.72 seconds for task-state data). By making this adjustment, we ensured that the resting-state DM coefficients, originally reflecting temporal evolution over 1.5 seconds, were scaled appropriately to match the temporal evolution over 0.72 seconds represented in the task-state DM coefficients.

### Optimizing the number of DMs

To determine the optimal number of DMs for accurately describing brain-wide BOLD dynamics, we employed a 5-fold cross-validation approach that evaluated predictive performance across models with increasing numbers of DMs. The number of DMs was incremented by sequentially including those with group-level DM evolution coefficients with large magnitudes (*λ*_*k*_; see “Group-level DMD” section), reflecting their relative contribution to the group-level dynamics. In each fold, the training set was used to estimate the group-level DMs (***ϕ***_*k*_). For the test set, each subject’s resting-state fMRI run was divided into two segments: (1) test-fitting data (first four-fifths, 12 minutes) — which used to estimate the subject-specific DM coefficients (*d*_*k*_); (2) test-validation data (remaining one-fifth, 3 minutes) — which was reserved for assessing predictive ability.

Predictive performance was evaluated by measuring the prediction error based on the entire history within the test-validation segment for each subject in the test set. The prediction error was defined as:

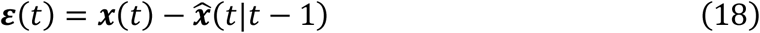

where ***x***(*t*) is the observed BOLD activation vector at time t, and 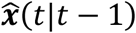 is the one-step-ahead prediction based on the DM coefficients fitted using the test-fitting data (see previous section).

For normalization, we utilized the *R*^2^ measure, which quantifies the proportion of variance in the signal that the model can explain. To account for varying variance across ROIs, we calculated *R*^2^ for each ROIs and averaged them:

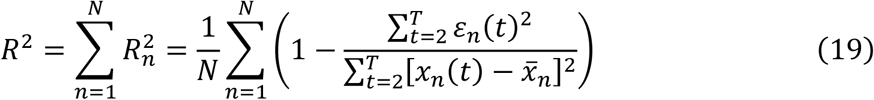

where *x*_*n*_(*t*) is the BOLD activation of the *n*-th ROI at time t, *ε*_*n*_(*t*) is the prediction error for the *n*-th ROI at time t, and 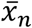 is the mean BOLD activation over time for the *n*-th ROI, calculated as: 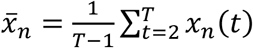.

This *R*^2^ measure was calculated for every subject in the test set of each fold, then averaged across the five folds, and subsequently used to compare model performance.

### Null model

To establish a baseline for model comparison, we used a null AR(1) model that accounts solely for the autocorrelation between temporally adjacent signals:

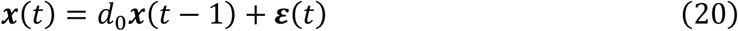

where *d*_0_ is the scalar autocorrelation coefficient and ε(*t*) represents white noise error terms. The optimal estimation of *d*_0_ is achieved using the following equation:

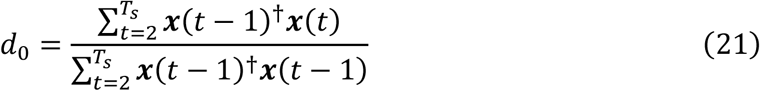

where ***x***(*t* − 1)^†^ represents the (complex conjugate) transpose of ***x***(*t* − 1).

Note that this null AR(1) model achieved a higher cross-validated *R*^2^ of 0.7767 compared to the zero-order random walk model (***x***(*t*) = ***x***(*t* − 1) + ε(*t*)), which achieved an *R*^2^ of 0.7618.

### Sparse linear model

In previous comparisons utilizing state-of-the-art models to characterize macroscopic BOLD dynamics [56], simple linear models consistently demonstrated the best fit for resting-state BOLD time series. These linear models capture the differences in BOLD signals between adjacent time points (*t* and *t* − 1) by employing a linear combination of signals from the preceding time point (*t* − 1). Building on the superior performance of the null AR(1) model, our approach models the residual variance not accounted for by the null AR(1) model through a linear combination of signals at the previous time point:

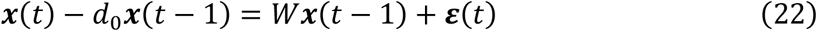

where *W* is an *N*-by-*N* matrix representing effective connectivity between ROIs, *d*_0_ is the scalar autocorrelation coefficient which is estimated identically to the null model, and ε(*t*) represents white noise error terms.

Due to the increased number of ROIs in our current study, estimating a dense linear model became infeasible. Thus, we employed a sparse modeling approach using LASSO regularization with a hyperparameter Γ. We estimated Γ by evaluating training data across a range of values [10,20,30, ⋯, 100] within each cross-validation fold. The optimal Γ was determined by interpolating the achieved cross-validated *R*^2^ values (estimated within the training dataset) across this range (with cubic spline interpolation) and subsequently applied to the test data (see Supplementary Figure 2a).

### Nonlinear manifold-based model

In the previous study [56], the manifold-based model achieved comparable fit for BOLD time series data among non-linear models. The manifold-based model uses the characteristics of manifold that can be approximated by a linear hyperplane in the small vicinity of a given point. We used the local polynomial modelling of order 1 [96] that is used in the previous work [56] but also with modification to model the residual variance not accounted for by the null AR(1) model:

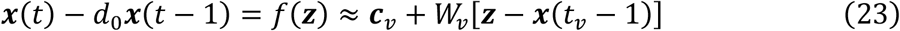

where the function *f*(***z***) is approximated as a linear function in the vicinity of ***x***(*t*_*l*_ − 1). The constant vector ***c***_*v*_ and matrix *W*_*v*_ were estimated from the test-fitting data ***x***(*t*_*f*_ − 1), but weighted according to its distance to ***x***(*t*_*v*_ − 1) with a gaussian kernel:

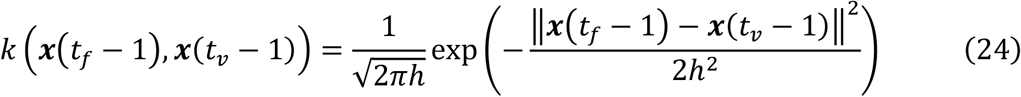

The hyperparameter *h* controls the degree of locality in the model. To ensure that the degree of locality in the model is appropriately scaled, the hyperparameter *h* is normalized based on the median of the distances between test-fitting and test-validation data (*h* · median(‖***x***(*t*_*f*_ − 1) − ***x***(*t*_*v*_ − 1)‖)/*h*_base_), where *h*_base_ = 50 (see the MATLAB code provided by Nozari, Bertolero [56]). We estimated *h* by evaluating training data across a range of values [2,4,6, ⋯, 20] within each cross-validation fold. The optimal *h* was determined by interpolating the achieved cross-validated *R*^2^ values (estimated within the training dataset) across this range (with cubic spline interpolation) and subsequently applied to the test data (see Supplementary Figure 2b).

For detailed methodology on the weighted least squares estimation used to solve this model, please refer to Nozari, Bertolero [56].

### Group-level FC topography

FC was determined by calculating Pearson correlations from each subject’s preprocessed ROI time-series without applying GSR. Specifically, FC was computed between 360 cortical regions for each subject, followed by Fisher’s r-to-z transformation (arctanh). These transformed FC values were then averaged across all subjects and converted back to correlation coefficients within the range of [−1, 1] using the tanh function.

### TVFC measure (std)

To assess the temporal variability of FC, we computed the std of TVFC using a sliding window method with a window size of *w* = 63 TRs (45.36 seconds). Prior to applying the sliding window approach, we performed high-pass filtering to remove frequency components below 1/*w* (where *w* is the window size in seconds) in addition to the previously applied bandpass filtering (0.01–0.1 Hz) to eliminate spurious fluctuations in TVFC [97]. Within each sliding window, Pearson correlation coefficients were computed for all pairs of cortical ROIs. The std of these TVFC time series for each ROI was then determined to quantify the temporal variability of FC.

### Reconstructed FC and TVFC matrices using DMs

To reconstruct the FC matrix, we aggregated contributions from the top five DMs. For each DM, we generated time series for all 360 cortical regions by normalizing the group-level DM evolution coefficients (*λ*_*k*_) to unit magnitude, thereby preserving only phase information, and applied these coefficients across 10,000 time points with a temporal resolution of 0.5 seconds. The time series for the *n*-the ROI in the *k*-th DM were reconstructed using the formula:

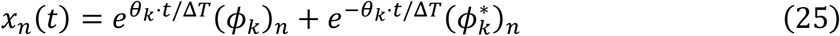

where (*ϕ*_*k*_)_*n*_ is the *n*-th element of the *k*-th DM vector *ϕ*_*k*_, *θ*_*k*_ is the phase (argument) of the group-level DM coefficient *λ*_*k*_, and Δ*T* is the resampled TR used in the group-level DMD (Δ*T* = 1.5*s*).

The five reconstructed time (i.e., one per each DM) series were then weighted and averaged based on the group-averaged mode engagement level of each 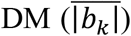. The FC matrix was subsequently computed by calculating Pearson correlation coefficients between all pairs of the averaged ROI time series. The TVFC (i.e., std) matrix was computed by calculating the std of FCs estimated using a sliding window approach, employing the same window size as used in the original TVFC analysis. To assess the similarity of the reconstruction, we correlated the lower triangular elements of the reconstructed FC matrix (or reconstructed TVFC matrix) with those of the original FC matrix (or original TVFC matrix).

### Network community analysis

For identify community structure, we applied the Newman’s spectral community detection implemented in Brain Connectivity Toolbox [98] with a resolution parameter set to 1 to both the original FC matrix and a reconstructed FC matrix derived from the DMs.

### FC gradients (cortical areas)

Following the methodology of the previous study [14], the group-level cortical FC matrix was thresholded to retain only the top 20% of values before extracting the gradients to consider most significant connections. Then, resting-state FC gradients were obtained by performing PCA on the group-level cortical FC matrix. See Figure 2b for topographies of the FC gradients.

### Subcortical and cerebellar FC-based functional gradients

Due to the lower signal-to-noise ratio of BOLD signals in non-neocortical regions compared to neocortical areas, we employed connectopic mapping [49] to characterize the functional organization of subcortical structures using resting-state fMRI data. This approach involves calculating the FC profiles between subcortical ROIs and other brain areas, followed by assessing the similarities of these FC profiles among subcortical ROIs. Subsequently, a dimensionality reduction technique is applied to the similarity matrix to derive spatial topographies that account for the most variance.

We applied this methodology to obtain the functional gradients of the hippocampus, amygdala, thalamus, striatum, brain stem, and cerebellum. For each subcortical area, let there be *N* ROIs within the area and *M* ROIs outside of it. For each subject, we computed the group-level FC matrix between the subcortical ROIs and external ROIs and then averaged these matrices across subjects to obtain a group-level FC topography. This process resulted in an *M*-by-*N* FC matrix, where each column represents the FC profile of a subcortical ROI. We then calculated the pairwise Pearson correlations between FC profiles of all subcortical ROIs, generating an *N*-by-*N* similarity (i.e., correlation) matrix. PCA was performed on this similarity matrix, and the first principal component, representing the principal functional gradient of the subcortical area, was retained. The same methodology was applied to the cerebellum to derive its functional gradient.

The resulting spatial topographies of the functional gradients for the subcortical areas and the cerebellum are illustrated in Supplementary Figure 6.

### Lag projection

We employed the lag-analysis MATLAB script provided by Raut, Mitra [99] on all cortical ROI time-series. Specifically, the time-lag between each pair of cortical ROI time series was determined by identifying the peak of their lagged cross-covariance function [51]. To achieve finer temporal resolution, the cross-covariance functions—originally calculated at a sampling rate of TR = 0.72s—were interpolated using three-point parabolic interpolation. A maximum shift of ±6 TR (±4.32s) was allowed when estimating the cross-covariance functions, and time lags longer than 4 s were excluded due to their high likelihood of being influenced by error or artifacts [99]. The lag projections were then calculated by averaging the pair-wise time-lag values across each column of the resulting time-lag matrix for all cortical ROIs.

### Quasi-periodic pattern (QPP)

QPPs were identified using the template matching algorithm described by Majeed, Magnuson [100]. The cortical ROI time-series from each subject were concatenated, and a random window of BOLD timepoints was selected as the initial QPP template. The algorithm then calculated the sliding correlation between the concatenated ROI time-series and this template. During the first two iterations, a correlation threshold of 0.1 was applied, which was increased to 0.2 for subsequent iterations. Peaks in the thresholded correlation time series were detected, and the corresponding time-series segments were averaged to update the QPP template. This iterative process was repeated until the template remained unchanged for two consecutive iterations (r > 0.9999). The algorithm was executed ten times, and the template with the highest average sliding window correlation across all iterations was selected as the final QPP template. QPP identification was performed on time series with and without GSR. GSR was performed by regressing the global mean time series—calculated as the average across all cortical ROIs—onto each individual cortical time series.

### Correlation between the global signal and the projected signal onto DMs

The global signal time series was derived by averaging the time series of all ROIs. To obtain the global activity time series of the *k*-th DM, we first calculated the coefficient *b*_*k*_(*t* − 1) for each time point (see “Subject-level estimation of DM metrics” section) and multiplied it by the group-level DM vector ***ϕ***_*k*_ (i.e., *b*_*k*_(*t* − 1)***ϕ***_*k*_). We then averaged the real components of these products across all ROIs to generate the global activity time series for the projected signal corresponding to the *k*-th DM. Specifically, the global activity of the *k*-th DM at time *t* – 1 calculated as 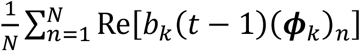 where *N* denotes the number of ROIs, and (***ϕ*** *k*)_*n*_ denotes the *n*-th element of DM vector ***ϕ***_*k*_. Finally, we computed the correlation between the global signal time series and the global activity time series of the projected signal for each subject.

### Seed-based FC

FC was calculated using ROI time series with the application of GSR. GSR was performed by regressing the global mean time series—calculated as the average across all cortical ROIs—onto each individual cortical time series. The residuals from these regressions were then utilized for determining FCs. FCs were determined by calculating Pearson correlations between the residual time series of each seed ROI and those of the other cortical ROIs.

### Yeo’s 7-network partition

The 7-network parcellation of the cortical areas was performed following Yeo, Krienen [17]. For this analysis, we used the group FC matrix (within cortical areas) with GSR. Each ROI’s connectivity profile was z-score normalized across all ROIs to ensure standardized data. K-means clustering was then applied to the normalized connectivity profiles, specifying seven clusters with 10 replicates to enhance clustering stability. The resulting clusters corresponded to two DMNs (possibly reflecting sub-networks of DMN), the CEN, SM, visual, dorsal attention, and ventral attention networks. The cluster centroids were utilized to represent the spatial distribution of each respective network. Supplementary Figure 4 illustrates the spatial topographies of each cluster centroid.

### Laterality index (LI)

To evaluate the hemispheric lateralization throughout the scanning period, we calculated the LI for each ROI. Specifically, we used the mean laterality index (MLI), which is the average of the dynamic laterality index (DLI) values across all sliding time windows as defined by Wu, Kong [52]. The procedure is outlined below.

Initially, we calculated the DLI for each ROI within sliding time windows of *w* = 30 seconds (~41.6TRs) with a 10-second sliding interval (Δ*w*). The DLI was defined using a global signal-based approach. Specifically, for the *n*-th ROI during the time window starting at time *t*, the DLI was computed as the difference between its Pearson correlation with the left hemispheric global signal (GS_L_) and its correlation with the right hemispheric global signal (GS_R_):

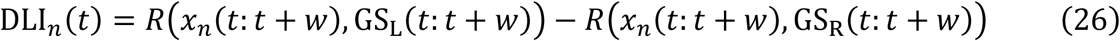

where *x*_*n*_(*t*: *t* + *w*) represents the BOLD time series of the *n*-th ROI within the time window from *t* to *t* + *w*. GS_L_(*t*: *t* + *w*) and GS_R_(*t*: *t* + *w*) denote the global signals of the left and right hemispheres, respectively, within the same time window. These are calculated as the averaged BOLD time series of all ROIs within each hemisphere. *R*(·,·) denotes the Pearson correlation coefficient.

A positive DLI indicates that the ROI’s activity is more synchronized with the left hemisphere (leftward laterality), while a negative DLI indicates stronger synchronization with the right hemisphere (rightward laterality).

The MLI for each ROI was derived by averaging its DLI values across all sliding time windows:

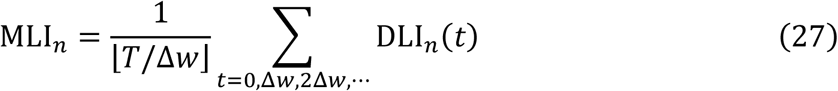

where *T* is the total number of time points, ⌊·⌋ is the floor function, ⌊*T*/Δ*w*⌋ is the total number of sliding time windows, DLI_*n*_(*t*) is the DLI of *n*-th ROI during the time window starting at time *t*. Thus, MLI indicates each ROI’s average hemispheric lateralization over the entire scan duration.

### Behavior analysis

For our factor analysis, we utilized the Python script provided by Schöttner, Bolton [64]. This script conducts hierarchical factor analysis on the dataset to identify underlying latent variables, employing maximum likelihood estimation in conjunction with orthogonal (varimax) rotation.

Running this script generates factor analysis results across various numbers of latent dimensions. We opted for a four-factor model due to its previously demonstrated robustness [64].

The four factors derived from the HCP dataset represent distinct dimensions of behavior. (1) Mental health reflects a spectrum of emotional well-being, encompassing positive states like life satisfaction and social support alongside challenges such as anxiety and depression. (2) Cognition captures intellectual abilities across various domains, including memory, reasoning, and language, indicating overall cognitive function. (3) Processing speed measures the efficiency of cognitive processing, emphasizing quick responses and reaction times, which contribute to cognitive performance in diverse tasks. (4) Substance use represents tendencies toward alcohol, tobacco, and drug use, highlighting behavioral patterns that can influence physical and mental health.

To examine the relationships between these behavioral factors and DM metrics while controlling for potential confounds, we performed general linear model (GLM) analyses using an ANCOVA approach. The confounding variables included in the model were age, sex, handedness, and the cube roots of ICV and TGMV.

Given the multiple comparisons involved in assessing all possible combinations of behavioral factors (four latent factors), DM metrics (five metrics), and metric types (three types: engagement, persistence, and progression), we applied a FWE correction to our statistical assessments. To achieve this, we employed Smith’s permutation testing method [101]. Specifically, all DM metrics were projected onto the vector space defined by the confounding variables and a constant term, retaining only the residuals. GLM analyses were then conducted using these residuals. To calculate FWE-corrected p-values, the residuals were permuted across subjects. In each permutation, GLM analyses were performed for all possible pairs of behavioral factors, DM metrics, and metric types, and the maximum absolute t-value was retained. This process was repeated for 10,000 permutations to generate a distribution of maximum absolute t-values, which was used to determine the p-values.

### Heritability analysis

We utilized twin data from the HCP to assess the heritability of DMs. Participants lacking zygosity information or essential confound data (e.g., age, sex, structural information) were excluded, resulting in a sample of 998 individuals (female = 532, mean age = 28.71, std = 3.72), including 104 monozygotic (MZ) twin pairs, 97 dizygotic (DZ) twin pairs, and 110 singletons. Zygosity information was employed to fit an ACE model, which decomposes the total phenotypic variance into three components: genetic factors (A, representing heritability), shared environmental (C), and unique environmental factors (E). The APACE permutation inference system [63] was applied using each subject’s DM metrics (i.e., mode-engagement level, persistence rate, and progression rate). For each DM metric, we regressed out the effects of the following confounding: age, age squared, sex, the interaction between age and sex, the interaction between sex and age squared, body mass index (BMI), race, handedness, years of education, cube root of ICV and TGMV, mean frame-wise displacement. The ACE model was applied separately to the three DM metrics. Statistical significance (p-values) was determined using 5,000 permutations, and confidence intervals (CIs) were estimated through 1,000 bootstrap resamples.

We conducted cosine similarity analysis using the magnitude and phase measures after regressing out confounding variables.

### Assessment of the significance of DM

If a specific DM is not significantly present in the subject’s BOLD signal dynamics, the estimation of the argument (phase) of its corresponding DM coefficient *d*_*k*_ (i.e., progression rate) tends to be randomly distributed around zero. This randomness arises because, without a substantial presence of the DM in the subject’s neural activity, there is insufficient information to consistently determine the direction of signal propagation associated with that DM. In such cases, the phase estimates do not carry meaningful information about the temporal evolution of the neural activity patterns within the mode.

Considering this, to assess whether each DM is significantly present (for the task-state data in the current study), we performed a one-sample t-test for each DM to determine if the mean phase significantly differs from zero across the subjects. A significant deviation from zero would indicate that the phase estimates are consistently oriented in a particular direction, suggesting a significant presence of the DM in the subject’s brain activity.

## Data availability

The resting-state and task-state fMRI data analyzed in this study are publicly accessible through the HCP S1200 Release. You can obtain the data at https://www.humanconnectome.org/study/hcp-young-adult/document/1200-subjects-data-release. For researchers interested in accessing restricted data, such as information on zygosity and parental details, please contact the HCP Project Manager and follow the application procedures outlined by the HCP at (https://www.humanconnectome.org/study/hcp-young-adult/document/restricted-data-usage) to obtain the data.

## Code availability

All analyses in this study were conducted using MATLAB R2021a. The scripts and code supporting the study’s findings are available on GitHub and will be publicly accessible upon the paper’s publication.

## References

1. Liu, X. and J.H. Duyn, Time-varying functional network information extracted from brief instances of spontaneous brain activity. Proceedings of the National Academy of Sciences, 2013. 110(11): p. 4392–4397.

2. Vidaurre, D., S.M. Smith, and M.W. Woolrich, Brain network dynamics are hierarchically organized in time. Proceedings of the National Academy of Sciences, 2017. 114(48): p. 12827–12832.

3. Shine, J.M., O. Koyejo, and R.A. Poldrack, Temporal metastates are associated with differential patterns of time-resolved connectivity, network topology, and attention. Proceedings of the National Academy of Sciences, 2016. 113(35): p. 9888–9891.

4. Greene, A.S., et al., Why is everyone talking about brain state? Trends in Neurosciences, 2023. 46(7): p. 508–524.

5. Breakspear, M., Dynamic models of large-scale brain activity. Nature neuroscience, 2017. 20(3): p. 340–352.

6. John, Y.J., et al., It’s about time: Linking dynamical systems with human neuroimaging to understand the brain. Network Neuroscience, 2022. 6(4): p. 960–979.

7. Calhoun, V.D., et al., The chronnectome: time-varying connectivity networks as the next frontier in fMRI data discovery. Neuron, 2014. 84(2): p. 262–274.

8. Pillai, A.S. and V.K. Jirsa, Symmetry breaking in space-time hierarchies shapes brain dynamics and behavior. Neuron, 2017. 94(5): p. 1010–1026.

9. Foster, M. and D. Scheinost, Brain states as wave-like motifs. Trends in Cognitive Sciences, 2024.

10. Song, H., W.M. Shim, and M.D. Rosenberg, Large-scale neural dynamics in a shared low-dimensional state space reflect cognitive and attentional dynamics. Elife, 2023. 12: p. e85487.

11. Raut, R.V., et al., Global waves synchronize the brain’s functional systems with fluctuating arousal. Science advances, 2021. 7(30): p. eabf2709.

12. Gu, Y., et al., Brain activity fluctuations propagate as waves traversing the cortical hierarchy. Cerebral cortex, 2021. 31(9): p. 3986–4005.

13. Vézquez-Rodríguez, B., et al., Signal propagation via cortical hierarchies. Network neuroscience, 2020. 4(4): p. 1072–1090.

14. Margulies, D.S., et al., Situating the default-mode network along a principal gradient of macroscale cortical organization. Proceedings of the National Academy of Sciences, 2016. 113(44): p. 12574–12579.

15. Collins, C.E., et al., Neuron densities vary across and within cortical areas in primates. Proceedings of the National Academy of Sciences, 2010. 107(36): p. 15927–15932.

16. Glasser, M.F. and D.C. Van Essen, Mapping human cortical areas in vivo based on myelin content as revealed by T1-and T2-weighted MRI. Journal of neuroscience, 2011. 31(32): p. 11597–11616.

17. Yeo, B.T., et al., The organization of the human cerebral cortex estimated by intrinsic functional connectivity. Journal of neurophysiology, 2011.

18. Wei, W., et al., A function-based mapping of sensory integration along the cortical hierarchy. Communications Biology, 2024. 7(1): p. 1–14.

19. Jung, H., T.D. Wager, and R.M. Carter, Novel cognitive functions arise at the convergence of macroscale gradients. Journal of Cognitive Neuroscience, 2022. 34(3): p. 381–396.

20. Choi, I., J.-Y. Lee, and S.-H. Lee, Bottom-up and top-down modulation of multisensory integration. Current Opinion in Neurobiology, 2018. 52: p. 115–122.

21. Mesulam, M.-M., From sensation to cognition. Brain: a journal of neurology, 1998. 121(6): p. 1013–1052.

22. Friston, K., A theory of cortical responses. Philosophical transactions of the Royal Society B: Biological sciences, 2005. 360(1456): p. 815–836.

23. Raichle, M.E., et al., A default mode of brain function. Proceedings of the national academy of sciences, 2001. 98(2): p. 676–682.

24. Greicius, M.D., et al., Functional connectivity in the resting brain: a network analysis of the default mode hypothesis. Proceedings of the national academy of sciences, 2003. 100(1): p. 253–258.

25. Menon, V., 20 years of the default mode network: A review and synthesis. Neuron, 2023. 111(16): p. 2469–2487.

26. Rottschy, C., et al., Modelling neural correlates of working memory: a coordinate-based meta-analysis. Neuroimage, 2012. 60(1): p. 830–846.

27. Vincent, J.L., et al., Evidence for a frontoparietal control system revealed by intrinsic functional connectivity. Journal of neurophysiology, 2008. 100(6): p. 3328–3342.

28. Menon, V. and M. D’Esposito, The role of PFC networks in cognitive control and executive function. Neuropsychopharmacology, 2022. 47(1): p. 90–103.

29. Fox, M.D., et al., The human brain is intrinsically organized into dynamic, anticorrelated functional networks. Proceedings of the National Academy of Sciences, 2005. 102(27): p. 9673–9678.

30. Taghia, J., et al., Uncovering hidden brain state dynamics that regulate performance and decision-making during cognition. Nature communications, 2018. 9(1): p. 2505.

31. Abbas, A., et al., Quasi-periodic patterns contribute to functional connectivity in the brain. Neuroimage, 2019. 191: p. 193–204.

32. Yousefi, B., et al., Quasi-periodic patterns of intrinsic brain activity in individuals and their relationship to global signal. Neuroimage, 2018. 167: p. 297–308.

33. Beaty, R.E., et al., Default and executive network coupling supports creative idea production. Scientific reports, 2015. 5(1): p. 10964.

34. Fornito, A., et al., Competitive and cooperative dynamics of large-scale brain functional networks supporting recollection. Proceedings of the National Academy of Sciences, 2012. 109(31): p. 12788–12793.

35. Menon, V., Large-scale brain networks and psychopathology: a unifying triple network model. Trends in cognitive sciences, 2011. 15(10): p. 483–506.

36. Menon, V. and L.Q. Uddin, Saliency, switching, attention and control: a network model of insula function. Brain structure and function, 2010. 214: p. 655–667.

37. Sridharan, D., D.J. Levitin, and V. Menon, A critical role for the right fronto-insular cortex in switching between central-executive and default-mode networks. Proceedings of the National Academy of Sciences, 2008. 105(34): p. 12569–12574.

38. Goulden, N., et al., The salience network is responsible for switching between the default mode network and the central executive network: replication from DCM. Neuroimage, 2014. 99: p. 180–190.

39. Seeley, W.W., The salience network: a neural system for perceiving and responding to homeostatic demands. Journal of Neuroscience, 2019. 39(50): p. 9878–9882.

40. Tseng, J. and J. Poppenk, Brain meta-state transitions demarcate thoughts across task contexts exposing the mental noise of trait neuroticism. Nature communications, 2020. 11(1): p. 1–12.

41. Vidaurre, D., et al., Behavioural relevance of spontaneous, transient brain network interactions in fMRI. Neuroimage, 2021. 229: p. 117713.

42. Bassett, D.S., et al., Dynamic reconfiguration of human brain networks during learning. Proceedings of the National Academy of Sciences, 2011. 108(18): p. 7641–7646.

43. Jin, C., et al., Dynamic brain connectivity is a better predictor of PTSD than static connectivity. Human brain mapping, 2017. 38(9): p. 4479–4496.

44. Kutz, J.N., et al., Dynamic mode decomposition: data-driven modeling of complex systems. 2016: SIAM.

45. Iungo, G.V., et al. Data-driven reduced order model for prediction of wind turbine wakes. in Journal of Physics: Conference Series. 2015. IOP Publishing.

46. Kuttichira, D.P., et al. Stock price prediction using dynamic mode decomposition. in 2017 International Conference on Advances in Computing, Communications and Informatics (ICACCI). 2017. IEEE.

47. Lu, H. and D.M. Tartakovsky, Prediction accuracy of dynamic mode decomposition. SIAM Journal on Scientific Computing, 2020. 42(3): p. A1639–A1662.

48. Barocio, E., et al., A dynamic mode decomposition framework for global power system oscillation analysis. IEEE Transactions on Power Systems, 2014. 30(6): p. 2902–2912.

49. Haak, K.V., A.F. Marquand, and C.F. Beckmann, Connectopic mapping with resting-state fMRI. Neuroimage, 2018. 170: p. 83–94.

50. Liu, T.T., A. Nalci, and M. Falahpour, The global signal in fMRI: Nuisance or Information? Neuroimage, 2017. 150: p. 213–229.

51. Mitra, A., et al., Lag structure in resting-state fMRI. Journal of neurophysiology, 2014. 111(11): p. 2374–2391.

52. Wu, X., et al., Dynamic changes in brain lateralization correlate with human cognitive performance. PLoS biology, 2022. 20(3): p. e3001560.

53. Glasser, M.F., et al., A multi-modal parcellation of human cerebral cortex. Nature, 2016. 536(7615): p. 171–178.

54. Ji, J.L., et al., Mapping the human brain’s cortical-subcortical functional network organization. Neuroimage, 2019. 185: p. 35–57.

55. Dawson, S.T., et al., Characterizing and correcting for the effect of sensor noise in the dynamic mode decomposition. Experiments in Fluids, 2016. 57: p. 1–19.

56. Nozari, E., et al., Macroscopic resting-state brain dynamics are best described by linear models. Nature biomedical engineering, 2024. 8(1): p. 68–84.

57. Luenberger, D.G., Introduction to Dynamic Systems: Theory, Models, and Applications. 1st ed. 1991: Wiley.

58. Cai, W., et al., Dynamic causal brain circuits during working memory and their functional controllability. Nature communications, 2021. 12(1): p. 3314.

59. Menon, V., et al., Optogenetic stimulation of anterior insular cortex neurons in male rats reveals causal mechanisms underlying suppression of the default mode network by the salience network. Nature Communications, 2023. 14(1): p. 866.

60. Fransson, P., Spontaneous low-frequency BOLD signal fluctuations: An fMRI investigation of the resting-state default mode of brain function hypothesis. Human brain mapping, 2005. 26(1): p. 15–29.

61. Gazzaley, A. and A.C. Nobre, Top-down modulation: bridging selective attention and working memory. Trends in cognitive sciences, 2012. 16(2): p. 129–135.

62. Gazzaley, A., et al., Functional interactions between prefrontal and visual association cortex contribute to top-down modulation of visual processing. Cerebral cortex, 2007. 17(Suppl_1): p. i125–i135.

63. Chen, X., E. Viding, and T. Nichols, Faster Accelerated Permutation Inference for the ACE Model (APACE) with Parallelization. 2017.

64. Schöttner, M., et al., Exploring the latent structure of behavior using the Human Connectome Project’s data. Scientific Reports, 2023. 13(1): p. 713.

65. Barch, D.M., et al., Function in the human connectome: task-fMRI and individual differences in behavior. Neuroimage, 2013. 80: p. 169–189.

66. Cole, M.W., et al., Intrinsic and task-evoked network architectures of the human brain. Neuron, 2014. 83(1): p. 238–251.

67. Bolt, T., et al., A parsimonious description of global functional brain organization in three spatiotemporal patterns. Nature Neuroscience, 2022. 25(8): p. 1093–1103.

68. Kelly, A.C., et al., Competition between functional brain networks mediates behavioral variability. Neuroimage, 2008. 39(1): p. 527–537.

69. Spreng, R.N., et al., Attenuated anticorrelation between the default and dorsal attention networks with aging: evidence from task and rest. Neurobiology of aging, 2016. 45: p. 149–160.

70. Hampson, M., et al., Functional connectivity between task-positive and task-negative brain areas and its relation to working memory performance. Magnetic resonance imaging, 2010. 28(8): p. 1051–1057.

71. Brincat, S.L., et al., Interhemispheric transfer of working memories. Neuron, 2021. 109(6): p. 1055–1066. e4.

72. Ringo, J.L., et al., Time is of the essence: a conjecture that hemispheric specialization arises from interhemispheric conduction delay. Cerebral Cortex, 1994. 4(4): p. 331–343.

73. Ocklenburg, S. and Z.V. Guo, Cross-hemispheric communication: Insights on lateralized brain functions. Neuron, 2024.

74. Thompson, G.J., et al., Short-time windows of correlation between large-scale functional brain networks predict vigilance intraindividually and interindividually. Human brain mapping, 2013. 34(12): p. 3280–3298.

75. Van den Heuvel, M.P. and O. Sporns, Network hubs in the human brain. Trends in cognitive sciences, 2013. 17(12): p. 683–696.

76. Gordon, E.M., et al., Three distinct sets of connector hubs integrate human brain function. Cell reports, 2018. 24(7): p. 1687–1695. e4.

77. Uddin, L.Q., et al., Functional connectivity of default mode network components: correlation, anticorrelation, and causality. Human brain mapping, 2009. 30(2): p. 625–637.

78. Lanzoni, L., et al., The role of default mode network in semantic cue integration. NeuroImage, 2020. 219: p. 117019.

79. Moneta, N., et al., Task state representations in vmPFC mediate relevant and irrelevant value signals and their behavioral influence. Nature Communications, 2023. 14(1): p. 3156.

80. O’Doherty, J., et al., Abstract reward and punishment representations in the human orbitofrontal cortex. Nature neuroscience, 2001. 4(1): p. 95–102.

81. Andrews-Hanna, J.R., et al., Functional-anatomic fractionation of the brain’s default network. Neuron, 2010. 65(4): p. 550–562.

82. Wang, H., et al., Evidence of a dissociation pattern in default mode subnetwork functional connectivity in schizophrenia. Scientific reports, 2015. 5(1): p. 14655.

83. Li, J., et al., Topography and behavioral relevance of the global signal in the human brain. Scientific reports, 2019. 9(1): p. 14286.

84. Murphy, K., et al., The impact of global signal regression on resting state correlations: are anti-correlated networks introduced? Neuroimage, 2009. 44(3): p. 893–905.

85. Murphy, K. and M.D. Fox, Towards a consensus regarding global signal regression for resting state functional connectivity MRI. Neuroimage, 2017. 154: p. 169–173.

86. Saad, Z.S., et al., Correcting brain-wide correlation differences in resting-state FMRI. Brain connectivity, 2013. 3(4): p. 339–352.

87. Deco, G., et al., The dynamic brain: from spiking neurons to neural masses and cortical fields. PLoS computational biology, 2008. 4(8): p. e1000092.

88. Jirsa, V.K. and H. Haken, Field theory of electromagnetic brain activity. Physical review letters, 1996. 77(5): p. 960.

89. Robinson, P.A., C.J. Rennie, and J.J. Wright, Propagation and stability of waves of electrical activity in the cerebral cortex. Physical Review E, 1997. 56(1): p. 826.

90. Van Essen, D.C., et al., The WU-Minn human connectome project: an overview. Neuroimage, 2013. 80: p. 62–79.

91. Smith, S.M., et al., Resting-state fMRI in the human connectome project. Neuroimage, 2013. 80: p. 144–168.

92. Bianciardi, M., et al., Sources of functional magnetic resonance imaging signal fluctuations in the human brain at rest: a 7 T study. Magnetic resonance imaging, 2009. 27(8): p. 1019–1029.

93. Cordes, D., et al., Frequencies contributing to functional connectivity in the cerebral cortex in “resting-state” data. American journal of neuroradiology, 2001. 22(7): p. 1326–1333.

94. Casorso, J., et al., Dynamic mode decomposition of resting-state and task fMRI. NeuroImage, 2019. 194: p. 42–54.

95. Biswal, B.B., et al., Toward discovery science of human brain function. Proceedings of the national academy of sciences, 2010. 107(10): p. 4734–4739.

96. Roll, J., Local and piecewise affine approaches to system identification. 2003: Univ.

97. Leonardi, N. and D. Van De Ville, On spurious and real fluctuations of dynamic functional connectivity during rest. Neuroimage, 2015. 104: p. 430–436.

98. Rubinov, M. and O. Sporns, Complex network measures of brain connectivity: uses and interpretations. Neuroimage, 2010. 52(3): p. 1059–1069.

99. Raut, R.V., et al., On time delay estimation and sampling error in resting-state fMRI. Neuroimage, 2019. 194: p. 211–227.

100. Majeed, W., et al., Spatiotemporal dynamics of low frequency BOLD fluctuations in rats and humans. Neuroimage, 2011. 54(2): p. 1140–1150.

101. Winkler, A.M., et al., Permutation inference for the general linear model. Neuroimage, 2014. 92: p. 381–397.

