## Supplementary Materials for "Large-scale Signal Propagation Modes in the Human Brain"

### Supplementary Results

*Supplementary Result 1 – Consistency of DMD results across cross-validation (CV) folds*

To determine the optimal number of DMs for characterizing the dynamics of brain-wide BOLD signals, we employed a five-fold CV approach as detailed in the main text. Group-level DMD was performed using the training data from each fold, consistently generating 12 DMs within the 0.01 Hz to 0.1 Hz frequency range per fold.

To evaluate the consistency of the DMD results across CV folds, resulting in a total of 60 DMs (i.e., 12 each for five folds), we calculated cosine similarity (CS) values for each DM pair. Initially, we applied the K-means clustering algorithm with K=12 to cluster similar DMs based on their magnitude vectors, where each element represents the magnitude of the corresponding DM vector component. This clustering accurately organized the DMs into 12 clusters, each containing 5 DMs, one from each of the five folds.

Subsequently, we assessed the similarity within each cluster (i.e., each containing 5 DMs, one from each CV fold) using cosine similarity. It is important to note that DM vectors remain equivalent despite any overall phase shifts in their elements (i.e., the signal propagations that they represent are invariant under multiplication by a complex scalar of unit magnitude), as phase is estimated at the subject level for each time point, thereby compensate for such shifts. However, standard CS for complex vectors is sensitive to these phase shifts. To address this, we utilized a phase-invariant cosine similarity (PICS), defined as the maximum CS obtained when one vector is arbitrarily phase-shifted relative to the other (defined by $e^{i\theta}$):

$$\mathrm{PICS}(\boldsymbol{\phi}_1, \boldsymbol{\phi}_2) = \max_{\theta \in [0,2\pi)} \left[\mathrm{CS}\left(\boldsymbol{\phi}_1, e^{i\theta}\boldsymbol{\phi}_2\right)\right] = \max_{\theta \in [0,2\pi)} \left[\frac{\mathrm{Re}\left(\boldsymbol{\phi}_1^{\dagger}\left(e^{i\theta}\boldsymbol{\phi}_2\right)\right)}{\|\boldsymbol{\phi}_1\|\|e^{i\theta}\boldsymbol{\phi}_2\|}\right] \quad \text{(S1)}$$

where † denotes conjugate transpose. Since $\left\|e^{i\theta}\boldsymbol{\phi}_2\right\| = \|\boldsymbol{\phi}_2\|$ and $\boldsymbol{\phi}_1^{\dagger}\left(e^{i\theta}\boldsymbol{\phi}_2\right) = e^{i\theta}\left(\boldsymbol{\phi}_1^{\dagger}\boldsymbol{\phi}_2\right)$, if we let $\boldsymbol{\phi}_1^{\dagger}\boldsymbol{\phi}_2 = re^{i\theta'}$, then $\boldsymbol{\phi}_1^{\dagger}\left(e^{i\theta}\boldsymbol{\phi}_2\right) = re^{i(\theta+\theta')}$, thus

$\max_{\theta\in[0,2\pi)}\left[\mathrm{Re}\left(\boldsymbol{\phi}_1^{\dagger}\left(e^{i\theta}\boldsymbol{\phi}_2\right)\right)\right] = r = \left|\boldsymbol{\phi}_1^{\dagger}\boldsymbol{\phi}_2\right|$. Therefore,

$$\mathrm{PICS}(\boldsymbol{\phi}_1, \boldsymbol{\phi}_2) = \frac{\left|\boldsymbol{\phi}_1^{\dagger}\boldsymbol{\phi}_2\right|}{\|\boldsymbol{\phi}_1\|\|\boldsymbol{\phi}_2\|} \tag{S2}$$

Our analysis revealed that the DMs across CV folds exhibited high overall similarity, with PICS values exceeding 0.79. Therefore, the 12 DMs were highly consistent across cross-validation folds. Notably, the five DMs central to the current study demonstrated exceptional similarity, with PICS values greater than 0.96. Supplementary Figure 1 illustrates the PICS values for each DM.

*Supplementary Result 2 – Deriving the subject-level estimation of DM coefficients* $d_k$

We modeled the linear dynamics equation of the brain-wide BOLD signal for subject *s* is as following:

$$\boldsymbol{x}_s(t) = A_s\boldsymbol{x}_s(t-1) \tag{S3}$$

where $A_s \in \mathbb{R}^{N\times N}$ is the subject-specific connectivity matrix and $\boldsymbol{x}_s(t) \in \mathbb{R}^N$ represents the BOLD activation vector at time t for the *s*-th subject.

We have *N* ROIs in the brain. Since the group-level DMs are estimated, the connectivity matrix A will be approximated using these DMs:

$$A_s \approx d_0(I - \Phi\Phi^+) + \Phi D_s\Phi^+ \tag{S4}$$

where $\Phi \in \mathbb{C}^{N\times 2K}$ denote the complex matrix, whose columns are the DM vectors. $\Phi^+$ the pseudo-inverse of $\Phi$. *I* is a *N*-by-*N* identity matrix, and $I - \Phi\Phi^+$ indicate the residual matrix. $D_s$ is a *2K*-by-*2K* diagonal matrix containing the subject-specific evolution coefficients $d_k$ and their complex conjugates. $d_0$ represents the autocorrelation coefficient of the orthogonal complement of the column space of $\Phi$ (see the main text).

By introducing a set of parameters $\{d_0', d_1', \cdots, d_{2K}'\}$, we rewrite the approximation as:

$$A_s = d_0'(I - \Phi\Phi^+) + \Phi\left[\begin{pmatrix} d_1' & \cdots & 0 \\ \vdots & \ddots & \vdots \\ 0 & \cdots & d_{2K}' \end{pmatrix}\right]\Phi^+$$

$$= d_0'(I - SS^+) + \sum_{k=1}^{2K} d_k' \boldsymbol{u}_k \boldsymbol{v}_k^\dagger \qquad \text{(S5)}$$

where $\boldsymbol{u}_i$ is $i$-th column vector of $\Phi$, and $\boldsymbol{v}_i$ is $i$-th column vector of $(\Phi^+)^\dagger$. † denotes conjugate transpose. The variable $d_k'$ is an unknown complex parameter corresponding to the subject-specific evolution coefficient, while $d_0'$ is another unknown complex parameter representing the subject-specific autocorrelation coefficient of the residual.

For simplicity of derivation, we temporarily relax the requirement that the $d_k'$ values form pairs of complex conjugates and $d_0'$ is a real value. Despite this relaxation, accurate estimations of the evolution coefficient $d_k$, its complex conjugate $d_k^*$, and autocorrelation coefficient for residual $d_0$ are still achieved because $\Phi$ consists of pairs of complex conjugate vectors and $\boldsymbol{x}_s(t)$ is a real vector for all $t$=1,2,…,$T_s$.

The following derivation is based on the previous work by Casorso et al. (2019), with modifications to incorporate residual autocorrelation and to handle derivatives with respect to complex-valued parameters.

We want to minimize the following terms regarding all $d_k'$ for $k$=0,1, …, 2$K$:

$$F_s = \sum_{t=2}^{T_s} \|\boldsymbol{x}_s(t) - (d_0'(I - \Phi\Phi^+) + \Phi D_s \Phi^+)\boldsymbol{x}_s(t-1)\|^2 \qquad \text{(S6)}$$

This is a convex minimization problem. So, we calculated the condition of local minima using partial derivatives regarding $d_k'$ for $k$=0,1, …, 2$K$.

Define the error vector $\boldsymbol{e}(t)$:

$$\boldsymbol{e}(t) = (d_0'(I - \Phi\Phi^+) + \Phi D_s \Phi^+)\boldsymbol{x}_s(t-1) - \boldsymbol{x}_s(t) \qquad \text{(S7)}$$

Then,

$$F_s = \sum_{t=2}^{T_s} \boldsymbol{e}(t)^\dagger \boldsymbol{e}(t) \tag{S8}$$

Since $d_k'$ is a complex variable, $F_s$ should be minimized with respect to $x_k = \mathrm{Re}(d_k')$ and $y_k = \mathrm{Im}(d_k')$ for all $k$. This can be easily achieved by adapting Wirtinger derivatives, where the partial derivatives are defined as follows:

$$\frac{\partial}{\partial d_k'} = \frac{1}{2}\left(\frac{\partial}{\partial x_k} + \frac{1}{i}\frac{\partial}{\partial y_k}\right) \tag{S9a}$$

$$\frac{\partial}{\partial {d_k'}^*} = \frac{1}{2}\left(\frac{\partial}{\partial x_k} - \frac{1}{i}\frac{\partial}{\partial y_k}\right) \tag{S9b}$$

Getting back to our problem, according to Wirtinger derivatives, the partial derivatives for ${d_k'}^*$:

$$\frac{\partial F_s}{\partial {d_k'}^*} = \sum_{t=2}^{T_s} \left\{ \frac{\partial \boldsymbol{e}(t)^\dagger}{\partial {d_k'}^*} \boldsymbol{e}(t) \right\} \tag{S10}$$

Therefore,

$$\frac{\partial F_s}{\partial {d_0'}^*} = \sum_{t=2}^{T_s} \{\boldsymbol{x}_s(t-1)^\dagger (I - \Phi\Phi^+)^\dagger \boldsymbol{e}(t)\}$$

$$= \sum_{t=2}^{T_s} \{\boldsymbol{x}_s(t-1)^\dagger (I - \Phi\Phi^+)^\dagger [(d_0'(I - \Phi\Phi^+) + \Phi D_s \Phi^+)\boldsymbol{x}_s(t-1) - \boldsymbol{x}_s(t)]\}$$

$$= \sum_{t=2}^{T_s} \left\{ \boldsymbol{x}_s(t-1)^\dagger (I - \Phi\Phi^+)^\dagger d_0' (I - \Phi\Phi^+) \boldsymbol{x}_s(t-1) \right.$$

$$+ \boldsymbol{x}_s(t-1)^\dagger (I - \Phi\Phi^+)^\dagger \left( \sum_{k=1}^{2K} d_k' \boldsymbol{u}_k \boldsymbol{v}_k^\dagger \right) \boldsymbol{x}_s(t-1)$$

$$\left. - \boldsymbol{x}_s(t-1)^\dagger (I - \Phi\Phi^+)^\dagger \boldsymbol{x}_s(t) \right\}$$

(S11)

and

$$\frac{\partial F_s}{\partial {d'_k}^*} = \sum_{t=2}^{T_s} \{\boldsymbol{x}_s(t-1)^\dagger \boldsymbol{v}_k \boldsymbol{u}_k^\dagger \vec{e}(t)\}$$

$$= \sum_{t=2}^{T_s} \{\boldsymbol{x}_s(t-1)^\dagger \boldsymbol{v}_k \boldsymbol{u}_k^\dagger [d'_0(I - \Phi\Phi^+)\boldsymbol{x}_s(t-1) + \Phi D_s \Phi^+ \boldsymbol{x}_s(t-1) - \boldsymbol{x}_s(t)]\}$$

$$= \sum_{t=2}^{T_s} \left\{ \boldsymbol{x}_s(t-1)^\dagger \boldsymbol{v}_k \boldsymbol{u}_k^\dagger d'_0 (I - \Phi\Phi^+) \boldsymbol{x}_s(t-1) \right.$$

$$\left. + \boldsymbol{x}_s(t-1)^\dagger \boldsymbol{v}_k \boldsymbol{u}_k^\dagger \left( \sum_{k'=1}^{2K} d'_{k'} \boldsymbol{u}_{k'} \boldsymbol{v}_{k'}^\dagger \right) \boldsymbol{x}_s(t-1) - \boldsymbol{x}_s(t-1)^\dagger \boldsymbol{v}_k \boldsymbol{u}_k^\dagger \boldsymbol{x}_s(t) \right\} \tag{S12}$$

The local minima satisfy the following for all $k$:

$$\frac{\partial F_s}{\partial {d'_k}^*} = 0 \tag{S13}$$

These conditions result in a system of $2K+1$ linear equations with $2K+1$ unknown variables. Please note that $\frac{\partial F_s}{\partial d'_k} = 0$ condition gives the same (so redundant) equations since $F_s$ itself is a real value.

The coefficients of the linear equations can be expressed as a matrix like the following:

$$Z \begin{bmatrix} d'_0 \\ d'_1 \\ \vdots \\ d'_{2K} \end{bmatrix} = \boldsymbol{w} \tag{S14}$$

where $Z \in \mathbb{C}^{(2K+1)\times(2K+1)}$ is a square matrix, and $\boldsymbol{w} \in \mathbb{C}^{(2K+1)}$ is the right-hand side vector. The elements of $Z$ and $\boldsymbol{w}$ are like this $(i,j \in [1,2,\cdots,2K])$:

For $Z$:

$$z_{00} = \sum_{t=2}^{T_s} \boldsymbol{x}_s(t-1)^\dagger (I - \Phi\Phi^+)^\dagger (I - \Phi\Phi^+) \boldsymbol{x}_s(t-1) \tag{S15a}$$

$$z_{0i} = \sum_{t=2}^{T_s} \boldsymbol{x}_s(t-1)^\dagger (I - \Phi\Phi^+)^\dagger \, \boldsymbol{u}_i \boldsymbol{v}_i^\dagger \boldsymbol{x}_s(t-1) \tag{S15b}$$

$$z_{i0} = z_{0i}^\dagger \tag{S15c}$$

$$z_{ij} = \sum_{t=2}^{T_s} \boldsymbol{x}_s(t-1)^\dagger \left(\boldsymbol{v}_i \boldsymbol{u}_i^\dagger \boldsymbol{u}_j \boldsymbol{v}_j^\dagger\right) \boldsymbol{x}_s(t-1) \tag{S15d}$$

For *W*:

$$w_0 = \sum_{t=2}^{T_s} \boldsymbol{x}_s(t-1)^\dagger (I - \Phi\Phi^+)^\dagger \boldsymbol{x}_s(t) \tag{S16a}$$

$$w_i = \sum_{t=2}^{T_s} \boldsymbol{x}_s(t-1)^\dagger \boldsymbol{v}_i \boldsymbol{u}_i^\dagger \boldsymbol{x}_s(t) \tag{S16b}$$

By solving this linear system for each subject, we obtained the subject-specific DM coefficients:

$$\begin{bmatrix} d_0' \\ d_1' \\ \vdots \\ d_{2K}' \end{bmatrix} = Z^{-1} \boldsymbol{w} \tag{S17}$$

■

Note that the half of the $d_k'$ values are the complex conjugates of the other half, since $\Phi$ consists of pairs of complex conjugate vectors.

**Supplementary Figures**

*Supplementary Figure 1*

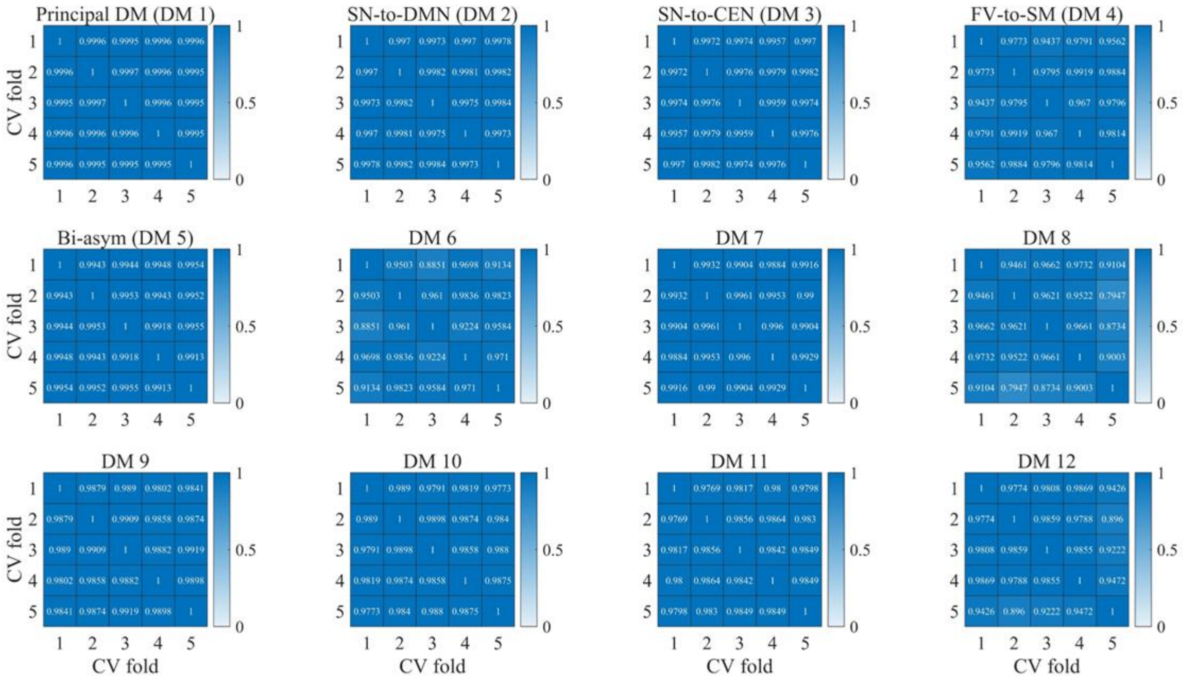


**Supplementary Figure 1.** Phase-Invariant Cosine Similarities (PICS) of each DM vector across five CV folds. Overall, all DMs exhibited high similarity (PICS > 0.79). Notably, except for DM 8 and DM 12, the similarities were exceptionally high (PICS > 0.96).

*Supplementary Figure 2*

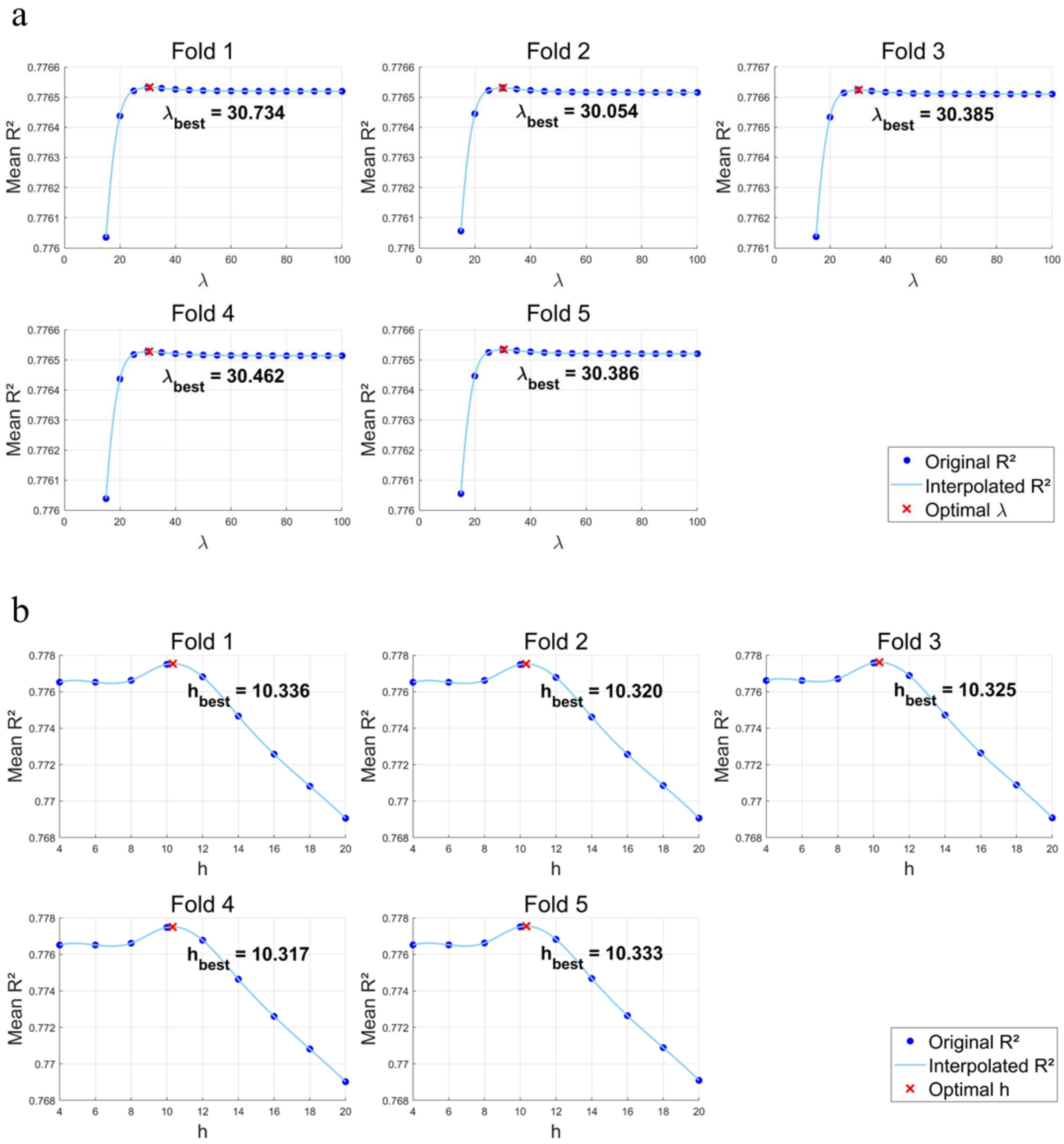


**Supplementary Figure 2.** Hyperparameter optimization for linear (LASSO) and nonlinear (manifold-based) fMRI dynamics models. Each model underwent a grid search (see Methods) to identify the hyperparameter configuration that maximized cross-validated $R^2$. For every candidate hyperparameter setting (indicated by blue dots), mean $R^2$ values were calculated from the training sets of each cross-validation fold. These results were then interpolated using cubic spline interpolation at finer hyperparameter increments (0.001). The optimal hyperparameter was selected based on the highest interpolated $R^2$ value, which was used for the test set. **(a)** Mean $R^2$ as a function of the hyperparameter for the LASSO model. **(b)** Mean $R^2$ as a function of the hyperparameter for the manifold-based model.

*Supplementary Figure 3*

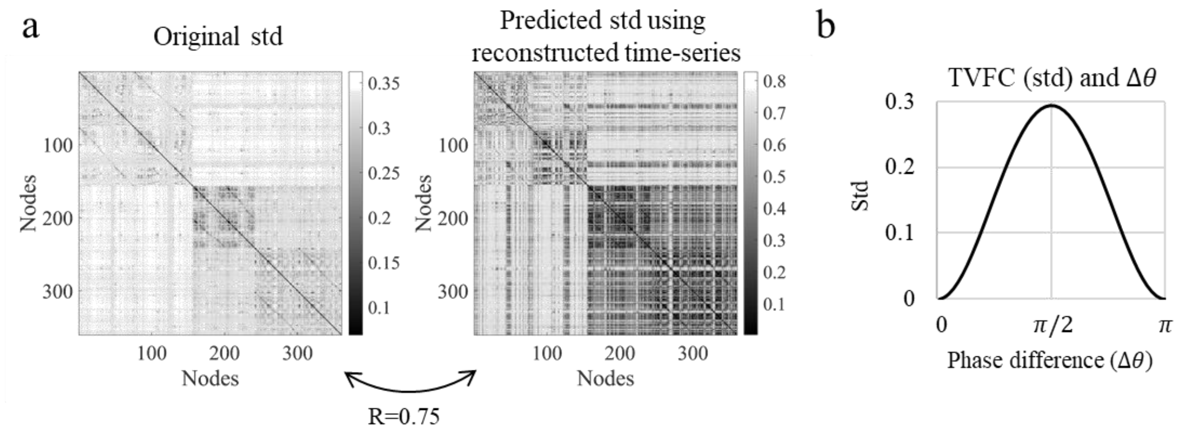


**Supplementary Figure 3**. Time-varying FC (TVFC) results. **(a) Left Panel**: Standard deviation (std) matrix of the original cortical BOLD time courses (i.e., the original std matrix). **Right Panel**: Std matrix reconstructed from the five DMs (refer to Methods for details). The std patterns between the original and reconstructed matrices exhibit a high degree of similarity, with a correlation coefficient of R = 0.75. **(b)** Illustration of the relationship between the std of temporal fluctuations in FC and the phase difference ($\Delta\theta$) between two sine waves. When the length of the sliding window used to estimate TVFC does not exactly match the period of the sine waves, the estimated FC fluctuates over time because the instantaneous relationship between the two sine waves varies. For mathematical details, see Leonardi and Van De Ville (2015). In this study, we generated two 1 Hz sine wave time series, each lasting 60 seconds with a sampling rate of 0.01 s, and varied the phase differences between them. We then estimated the correlation between the two sine waves using sliding windows of 0.7 s. The std is zero when the phase difference $\Delta\theta = 0$ or $\pi$, and it reaches its maximum at $\Delta\theta = \pi/2$.

*Supplementary Figure 4*

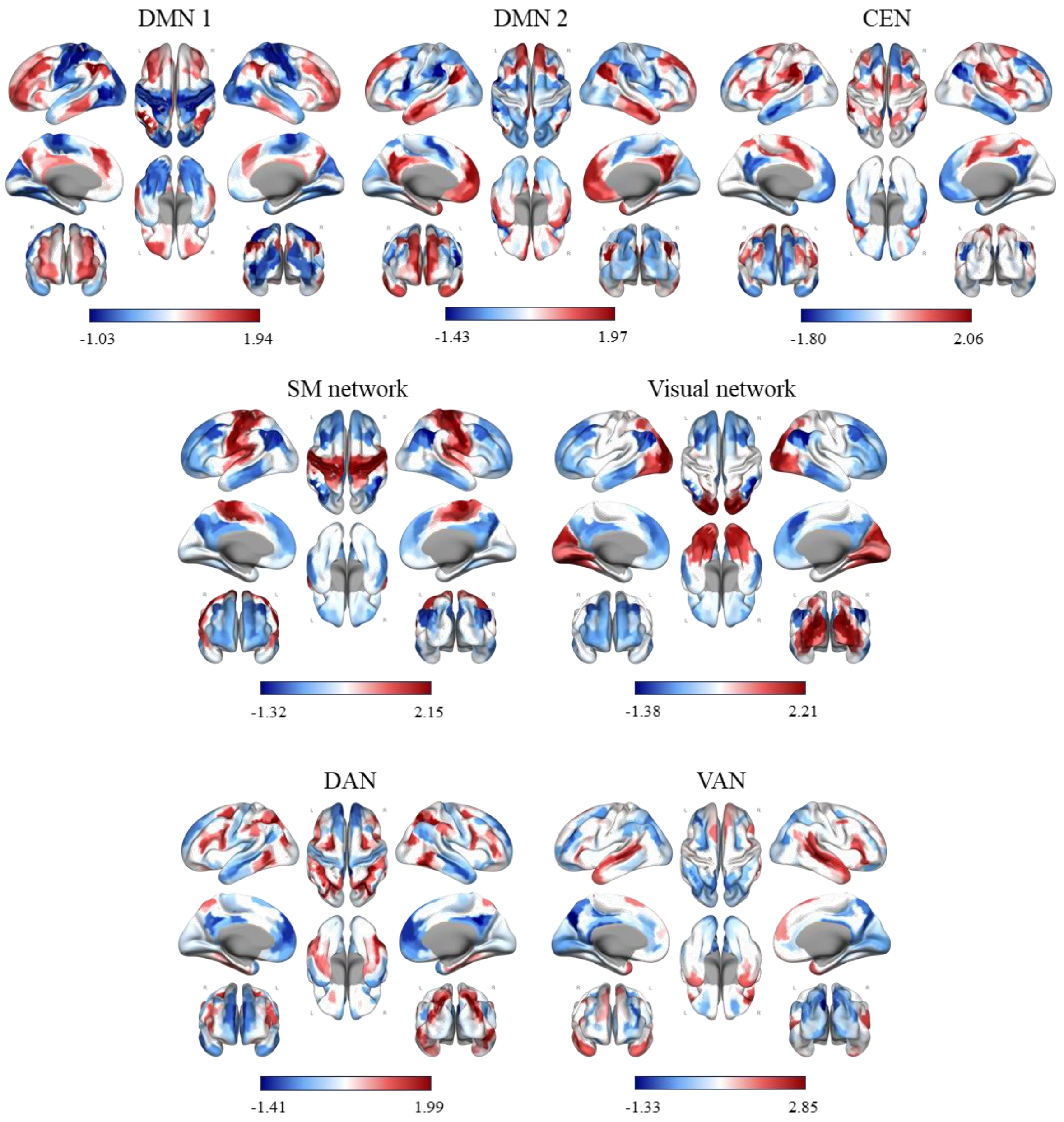


**Supplementary Figure 4.** Cluster centroids obtained from the clustering analysis of each ROI connectivity profile – Yeo's 7-network partition (Yeo et al., 2011). Network labels were assigned based on their spatial topographies. Abbreviations: DMN, default mode network; CEN, central executive network; SM network, sensorimotor network; DAN, dorsal attention network; VAN, ventral attention network.

*Supplementary Figure 5*

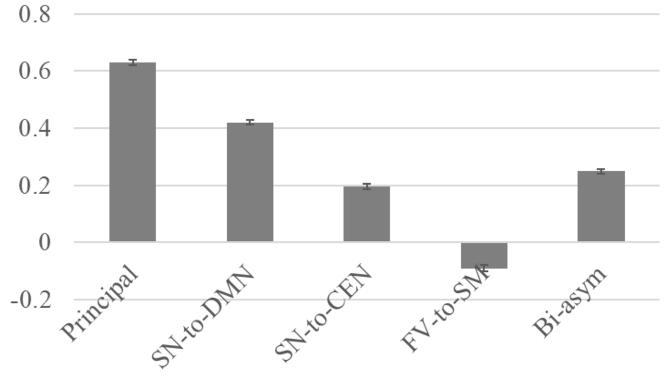


**Supplementary Figure 5.** The subject-averaged correlation between global signal and the global activity of projected fMRI signal onto the five DMs (see Methods). The error bars indicate 95% CI.

*Supplementary Figure 6*

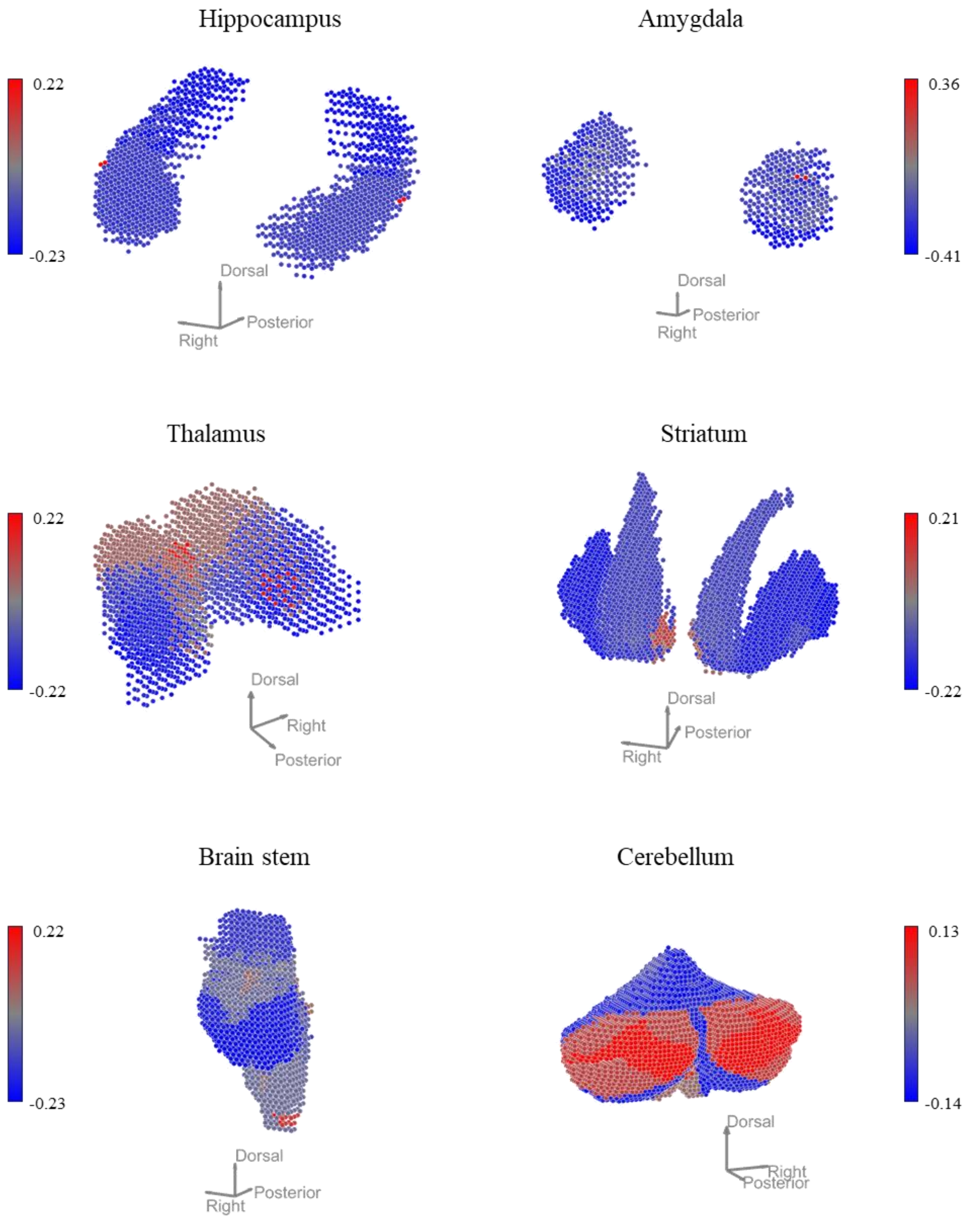


**Supplementary Figure 6.** Topographies of the FC-based functional gradient (the first PC) of the subcortical and cerebellar areas.

*Supplementary Figure 7*

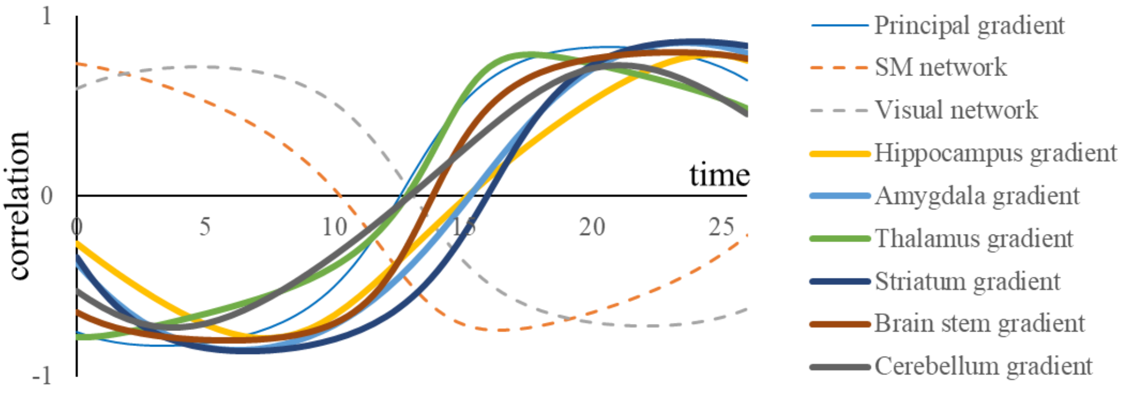


**Supplementary Figure 7.** Correlation between the activation of the principal DM and the functional gradients of subcortical and cerebellar regions. Activation time series were reconstructed using the principal DM, and correlations were calculated between the spatial pattern of activation at each time point and the functional gradients. (See Supplementary Figure 6 for the topographies of these subcortical and cerebellar functional gradients.) Additionally, correlations for the principal cortical gradient and the SM and visual networks from Yeo's 7-network partition are shown (identical to the inset of Figure 2a) for comparison. All correlations with the functional gradients exhibited similar variation patterns to those of the principal gradient, reaching peak values exceeding 0.7. See Supplementary Video 1 for the time course of the principal DM's brain-wide activation.

*Supplementary Figure 8*

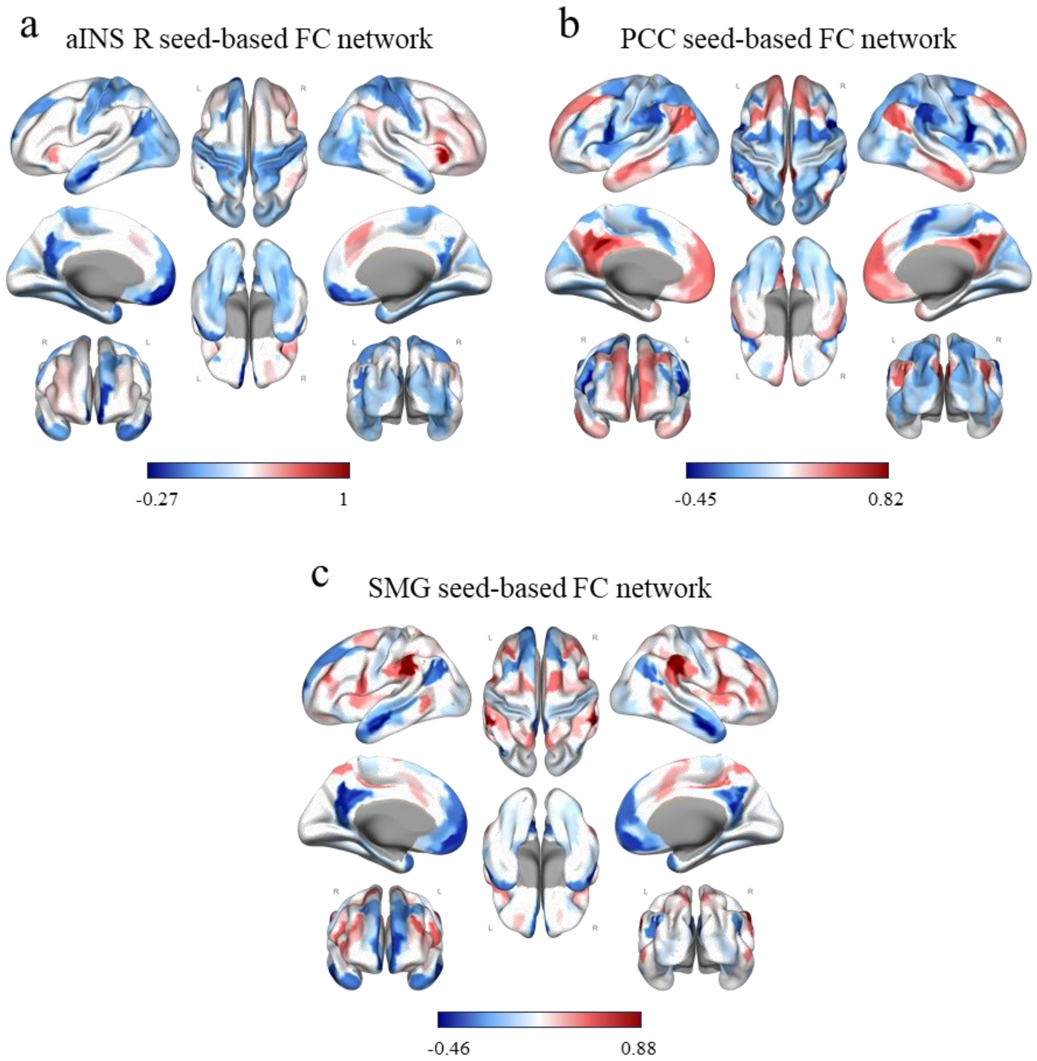


**Supplementary Figure 8.** Spatial distributions of seed-based FC networks. **(a)** FC network originating from the right anterior insula (aINS R) seed. **(b)** FC network originating from the posterior cingulate cortex (PCC) seed. **(c)** FC network originating from the supramarginal gyrus (SMG) seed.

*Supplementary Figure 9*

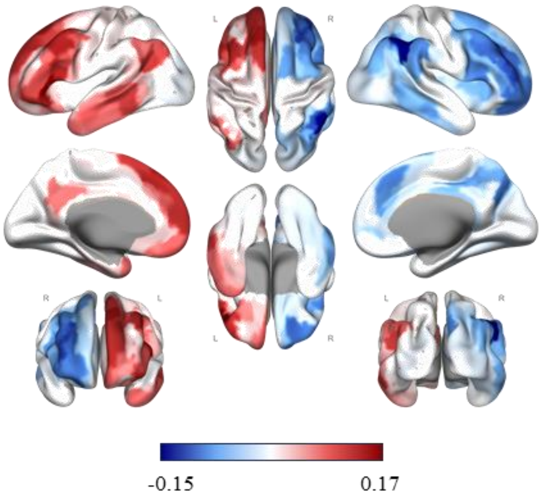


**Supplementary Figure 9.** Spatial distribution of the mean lateralization index (MLI). Positive MLI values indicate stronger synchronization with the left hemisphere (leftward laterality), whereas negative values indicate stronger synchronization with the right hemisphere (rightward laterality).

*Supplementary Figure 10*

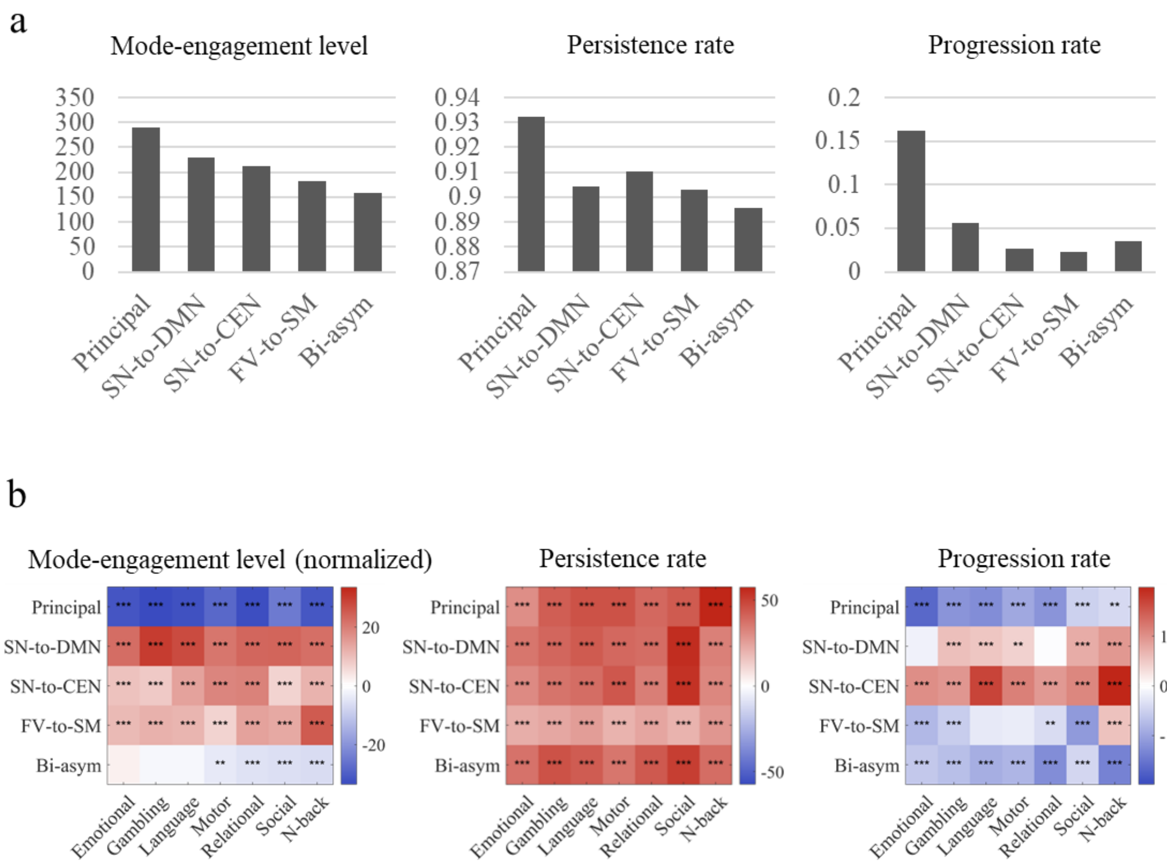


**Supplementary Figure 10. (a)** Average values of DM metrics across subjects. **(b)** T-value heatmaps comparing DM metrics between resting and task states. **Left panel:** Normalized mode-engagement rate. **Middle panel:** Persistence rates. **Right panel:** Progression rates. The left and right panels correspond to those shown in Figure 5d and 5e of the main text. P-values were Bonferroni-corrected separately for each of the three DM metric types (i.e., normalized mode-engagement rate, persistence rates, and progression rates).

*Supplementary Figure 11*

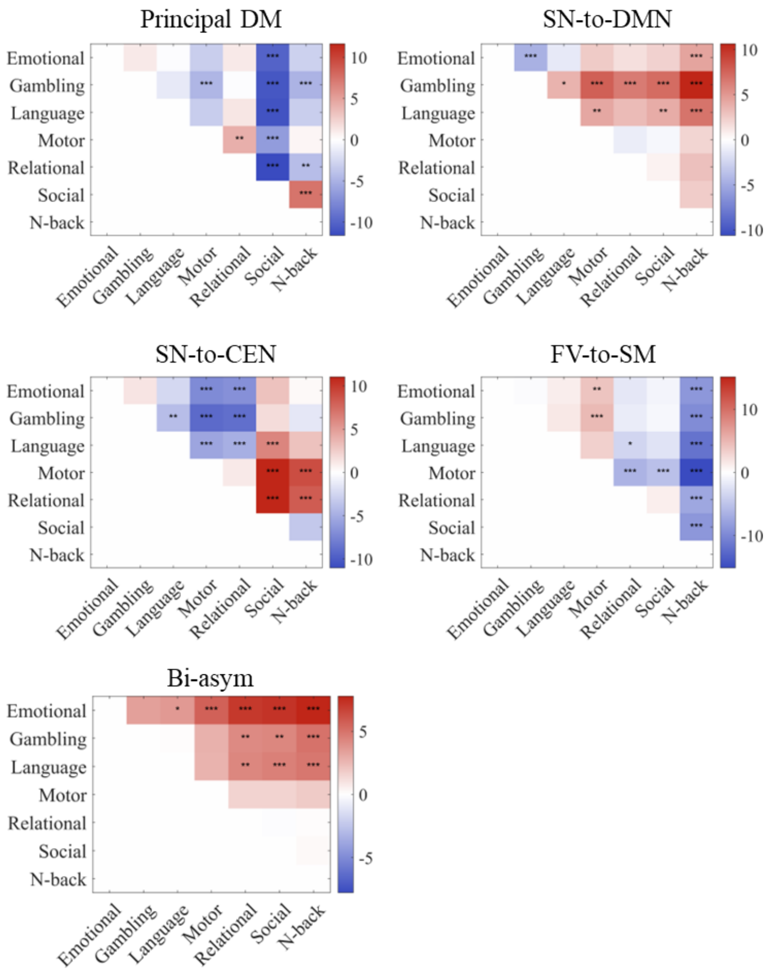


**Supplementary Figure 11.** T-value heatmaps comparing the normalized mode-engagement levels of the five DMs across all possible pairs of seven tasks. P-values for normalized mode-engagement levels were Bonferroni-corrected for the five DMs and all pairwise task comparisons. (*p < 0.05, **p < 0.01, ***p < 0.001)

*Supplementary Figure 12*

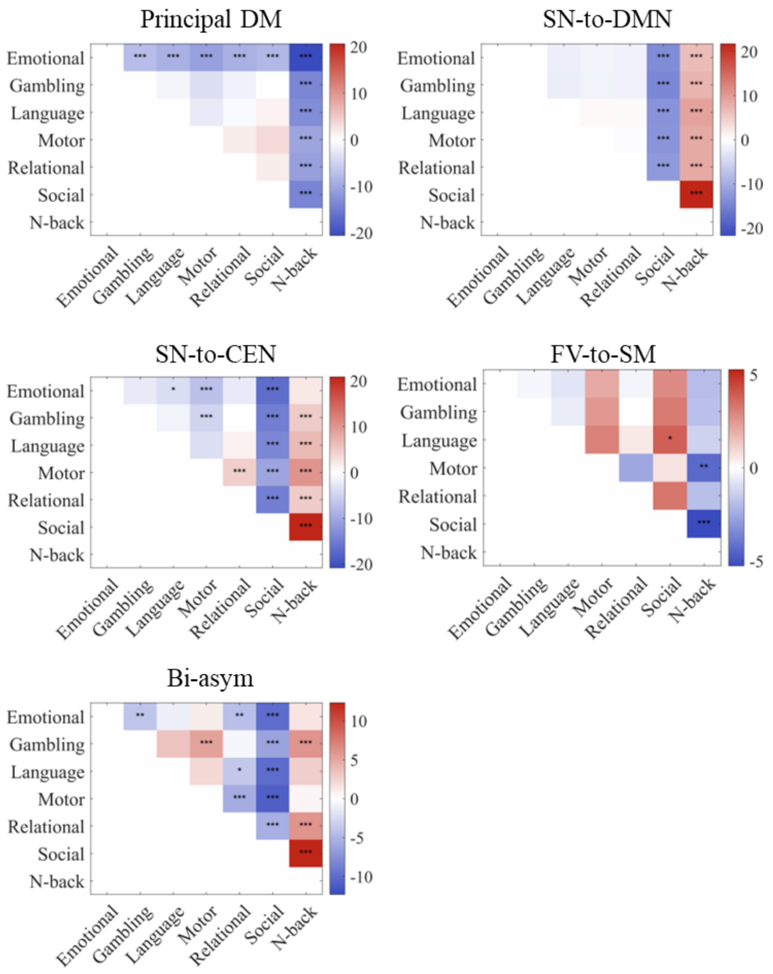


**Supplementary Figure 12.** T-value heatmaps comparing the persistence rates of the five DMs across all possible pairs of seven tasks. P-values for persistence rates were Bonferroni-corrected for the five DMs and all pairwise task comparisons. (*p < 0.05, **p < 0.01, ***p < 0.001)

*Supplementary Figure 13*

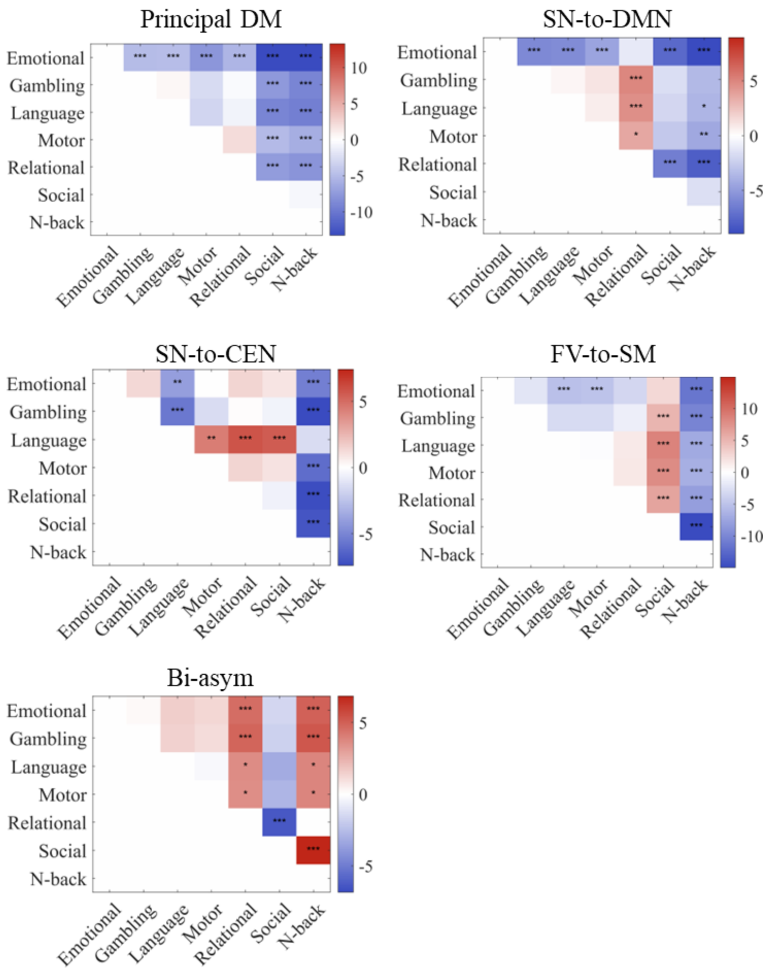


**Supplementary Figure 13.** T-value heatmaps comparing the progression rates of the five DMs across all possible pairs of seven tasks. P-values for progression rates were Bonferroni-corrected for the five DMs and all pairwise task comparisons. (*$p < 0.05$, **$p < 0.01$, ***$p < 0.001$)

**Supplementary Tables**

*Supplementary Table 1. Behavior results: t-values of mode-engagement level*

| DM | Mental Health | Cognition | Processing Speed | Substance Use |
|---|---|---|---|---|
| **Principal** | -1.287 | -1.4615 | -3.3858 | 0.9831 |
| **SN-to-DMN** | 0.0021 | 3.6924 | 2.138 | -0.6363 |
| **SN-to-CEN** | -0.55 | -1.1613 | 3.4203 | -2.8499 |
| **FV-to-SN** | 2.9094 | -3.5113 | -0.2242 | 1.2218 |
| **SN-to-CEN** | 1.1031 | 3.2081 | 3.0755 | -0.3521 |

*Supplementary Table 2. Behavior results: FWE-corrected p-values of mode-engagement level*

| DM | Mental Health | Cognition | Processing Speed | Substance Use |
|---|---|---|---|---|
| **Principal** | 1 | 0.9998 | 0.0434 | 1 |
| **SN-to-DMN** | 1 | 0.0148 | 0.8611 | 1 |
| **SN-to-CEN** | 1 | 1 | 0.0382 | 0.2279 |
| **FV-to-SN** | 0.1881 | 0.0285 | 1 | 1 |
| **SN-to-CEN** | 1 | 0.074 | 0.1155 | 1 |

*Supplementary Table 3. Behavior results: t-values of persistence rate*

| DM | Mental Health | Cognition | Processing Speed | Substance Use |
|---|---|---|---|---|
| **Principal** | 0.1865 | -0.232 | -2.5275 | -1.2574 |
| **SN-to-DMN** | -0.5419 | 1.5394 | 0.2425 | -1.1361 |
| **SN-to-CEN** | -1.9196 | 3.7665 | 0.882 | -1.1178 |
| **FV-to-SN** | 1.0347 | 0.8149 | -1.1603 | 2.1414 |
| **SN-to-CEN** | 0.3671 | 2.5268 | 1.0606 | -0.5841 |

*Supplementary Table 4. Behavior results: FWE-corrected p-values of persistence rate*

| DM | Mental Health | Cognition | Processing Speed | Substance Use |
|---|---|---|---|---|
| **Principal** | 1 | 1 | 0.4934 | 1 |
| **SN-to-DMN** | 1 | 0.9988 | 1 | 1 |
| **SN-to-CEN** | 0.9642 | 0.0117 | 1 | 1 |
| **FV-to-SN** | 1 | 1 | 1 | 0.8587 |
| **SN-to-CEN** | 1 | 0.4945 | 1 | 1 |

*Supplementary Table 5. Behavior results: t-values of progression rate*

| DM | Mental Health | Cognition | Processing Speed | Substance Use |
|---|---|---|---|---|
| **Principal** | -1.9818 | 0.4266 | -1.9173 | -0.0117 |
| **SN-to-DMN** | -0.3268 | 2.5238 | -0.51 | 0.4311 |
| **SN-to-CEN** | 2.7482 | 0.959 | 0.4004 | -0.8959 |
| **FV-to-SN** | -0.2982 | -0.6895 | -0.4961 | 0.8232 |
| **SN-to-CEN** | -0.3986 | 4.9439 | 3.3887 | -0.9304 |

*Supplementary Table 6. Behavior results: FWE-corrected p-values of progression rate*

| DM | Mental Health | Cognition | Processing Speed | Substance Use |
|---|---|---|---|---|
| **Principal** | 0.9432 | 1 | 0.9651 | 1 |
| **SN-to-DMN** | 1 | 0.4971 | 1 | 1 |
| **SN-to-CEN** | 0.2988 | 1 | 1 | 1 |
| **FV-to-SN** | 1 | 1 | 1 | 1 |
| **SN-to-CEN** | 1 | $<1\times10^{-4}$ | 0.0432 | 1 |

**Supplementary Videos**

**Supplementary Video 1.** Temporal reconstruction of the *principal DM*'s spatiotemporal patterns in cortical, subcortical, and cerebellar regions. Time points are uniformly spaced at $\Delta t = 0.5s$, spanning a full period of this DM. The cortical spatiotemporal pattern is identical to that depicted in Figure 2a. Red indicates positive activation, while blue indicates negative activation.

**Supplementary Video 2.** Temporal reconstruction of the *SN-to-DMN DM*'s spatiotemporal patterns in cortical, subcortical, and cerebellar regions. Time points are uniformly spaced at $\Delta t = 0.5s$, spanning a full period of this DM. The cortical spatiotemporal pattern is identical to that depicted in Figure 3a. Red indicates positive activation, while blue indicates negative activation.

**Supplementary Video 3.** Temporal reconstruction of the *SN-to-CEN DM*'s spatiotemporal patterns in cortical, subcortical, and cerebellar regions. Time points are uniformly spaced at $\Delta t = 0.5s$, spanning a full period of this DM. The cortical spatiotemporal pattern is identical to that depicted in Figure 3b. Red indicates positive activation, while blue indicates negative activation.

**Supplementary Video 4.** Temporal reconstruction of the *FV-to-SM DM*'s spatiotemporal patterns in cortical, subcortical, and cerebellar regions. Time points are uniformly spaced at $\Delta t = 0.5s$, spanning a full period of this DM. The cortical spatiotemporal pattern is identical to that depicted in Figure 44a. Red indicates positive activation, while blue indicates negative activation.

**Supplementary Video 5.** Temporal reconstruction of the *bi-asym DM*'s spatiotemporal patterns in cortical, subcortical, and cerebellar regions. Time points are uniformly spaced at $\Delta t = 0.5s$, spanning a full period of this DM. The cortical spatiotemporal pattern is identical to that depicted in Figure 4b. Red indicates positive activation, while blue indicates negative activation.

**Supplementary references**


CASORSO, J., KONG, X., CHI, W., VAN DE VILLE, D., YEO, B. T. & LIéGEOIS, R. 2019. Dynamic mode decomposition of resting-state and task fMRI. *NeuroImage,* 194**,** 42-54.

LEONARDI, N. & VAN DE VILLE, D. 2015. On spurious and real fluctuations of dynamic functional connectivity during rest. *Neuroimage,* 104**,** 430-436.

YEO, B. T., KRIENEN, F. M., SEPULCRE, J., SABUNCU, M. R., LASHKARI, D., HOLLINSHEAD, M., ROFFMAN, J. L., SMOLLER, J. W., ZöLLEI, L. & POLIMENI, J. R. 2011. The organization of the human cerebral cortex estimated by intrinsic functional connectivity. *Journal of neurophysiology*.
